# Integrative Modelling of Signalling Network Dynamics Identifies Cell Type-selective Therapeutic Strategies for FGFR4-driven Cancers

**DOI:** 10.1101/2021.11.03.467180

**Authors:** Sung-Young Shin, Nicole J Chew, Milad Ghomlaghi, Anderly C Chüeh, Yunhui Jeong, Lan K. Nguyen, Roger J Daly

**Affiliations:** Cancer Program, Biomedicine Discovery Institute, Monash University, Melbourne, VIC 3800, Australia; Department of Biochemistry and Molecular Biology, Monash University, Melbourne, VIC 3800, Australia

**Author notes:** These authors contributed equally to the work (Sung-Young Shin, Nicole J Chew).

**Keywords:** cancer cell resistance, combination therapy, network rewiring, signal transduction, computational mechanistic modelling, cancer type-specific treatments

## Abstract

Oncogenic FGFR4 signalling represents a potential therapeutic target in various cancer types, including triple negative breast cancer (TNBC) and hepatocellular carcinoma (HCC). However, resistance to FGFR4 single-agent therapy remains a major challenge, emphasizing the need for effective combinatorial treatments. Our study sought to develop a comprehensive computational model of FGFR4 signalling and provide network-level insights into resistance mechanisms driven by signalling dynamics. Our integrated approach, combining computational network modelling with experimental validation, uncovered potent AKT reactivation following FGFR4 targeting in the TNBC cell line MDA-MB-453. By systematically simulating the model to analyse the effects of co-targeting specific network nodes, we were able to predict, and subsequently confirm through experimental validation, the strong synergy of co-targeting FGFR4 and AKT or specific ErbB kinases, but not PI3K. Incorporating protein expression data from hundreds of cancer cell lines, we then adapted our model to diverse cellular contexts. This revealed that while AKT rebound is common, it is not a general phenomenon. ERK reactivation, for example, occurs in certain cell types, including the FGFR4-driven HCC cell line Hep3B, where there is a synergistic effect of co-targeting FGFR4 and MEK, but not AKT. In summary, our study offers key insights into drug-induced network remodelling and the role of protein expression heterogeneity in targeted therapy responses. We underscore the utility of computational network modelling for designing cell type-selective combination therapies and enhancing precision cancer treatment.

**Significance:** This study underscores the potential of computational predictive modelling in deciphering mechanisms of cancer cell resistance to targeted therapies and in designing more effective, cancer type-specific combination treatments.

## Introduction

Aberrant signalling by specific members of the FGFR family, comprising FGFR1-4, occurs in a variety of human cancers, reflecting FGFR gene alterations and/or enhanced ligand expression (1). This has led to the development of selective small molecule drugs targeting these receptors, including Erdafitinib (an inhibitor of FGFR1-4), which was recently FDA-approved for patients with metastatic urothelial carcinoma (2). While initial interest focused on FGFR1-3, oncogenic roles for FGFR4 in a variety of cancers have now become evident (3). In breast cancer, enhanced FGFR4 expression and mutation is associated with metastatic progression, particularly in the lobular subtype (4), and high FGFR4 expression positively correlates with endocrine resistance (4,5). FGFR4 also drives phenotypic switching of luminal A breast cancers to a HER2-enriched gene expression phenotype (6), and is overexpressed in approximately one third of triple negative breast cancers (TNBCs) (7). A second cancer where FGFR4 is strongly implicated is hepatocellular carcinoma (HCC). Here, FGFR4 is activated as a consequence of overexpression of its ligand, FGF19, which is a ‘driver’ oncogene on the chromosome 11q13.3 amplicon (8,9). These oncogenic roles for FGFR4 have led to the development of BLU9931, BLU-554 (fisogatinib) and H3B-6527, small molecule drugs that selectively target FGFR4 and exhibit promising pre-clinical activity in FGF19-driven HCC (9–11). Both fisogatinib and H3B-6527 are currently under evaluation in clinical trials for patients with advanced HCC (e.g. NCT04194801 and NCT02834780).

While targeted therapies directed towards specific oncogenic protein kinases have greatly improved clinical management of particular cancers (12), intrinsic and acquired drug resistance remain a major problem (13). One mechanism that can limit the efficacy of targeted therapy is dynamic, adaptive rewiring of the cellular signalling network in response to treatment (1). For example, treatment with AKT or MEK inhibitors can lead to increased expression and/or activation of a suite of receptor tyrosine kinases (RTKs) that dampens the cellular response to the drug (14,15). Importantly, characterization of network adaptation in response to drug treatment can inform the rational design of combination therapies that exhibit improved efficacy (14).

Despite these advances, the complexity of kinase signalling networks presents a major roadblock to understanding mechanisms of adaptive resistance to targeted therapy (1). This complexity reflects the presence of features such as pathway crosstalk and both positive and negative feedback loops, and results in non-intuitive behaviour that makes logical prediction of signalling and biological outputs challenging. A powerful approach to address this issue is to integrate mathematical network modelling with experimental analysis in order to generate and validate predictive models. This can provide fundamental insights into the wiring of signalling networks and how this generates specific signalling behaviours and biological outputs (16,17). In addition, it can be exploited to identify novel therapeutic strategies, as exemplified by accurate prediction of synergistic drug combinations by establishing a quantitative mechanistic model of the EGFR-PYK2-MET signalling network in TNBC (18).

In this study, we developed a comprehensive mechanistic model of the FGFR4 signalling network and used an integrative approach of computational modelling and experimental studies to analyse how the network responds to anti-FGFR4 inhibition across different cancer types and to identify combinatorial approaches that circumvent drug resistance. Overall, our work establishes an integrative framework for network-level analysis of drug-induced signalling adaptation that can be applied not only to FGFR4 inhibition but also other kinase-directed targeted therapies.

## RESULTS

### FGFR4 inhibition rewires signalling networks in MDA-MB-453 cells leading to upregulation of AKT activity

To determine the effects of the selective and irreversible FGFR4 inhibitor BLU9931 on FGFR4 downstream signalling and cell proliferation in TNBC, the MDA-MB-453 cell line that exhibits an activating mutation in FGFR4 was utilised (19). The MDA-MB-468 cell line with no detected FGFRs was used as a negative control (20). MDA-MB-453 cells were treated with different concentrations of BLU9931 for 1 h. Total FGFR4 expression was not affected by BLU9931, but the treatment decreased pFGFR at concentrations of 3nM and higher (Fig. 1A), and this effect was paralleled by a significant decrease in downstream signalling proteins pFRS2, pERK and pAKT (Fig. 1A, S1). MDA-MB-453 cell proliferation was significantly decreased by BLU9931 at concentrations of 10nM or higher, starting from day 3 after treatment (Fig. 1B). However, BLU9931 did not have any effect on MDA-MB-468 cell proliferation (Fig. 1C).

**Figure 1.**
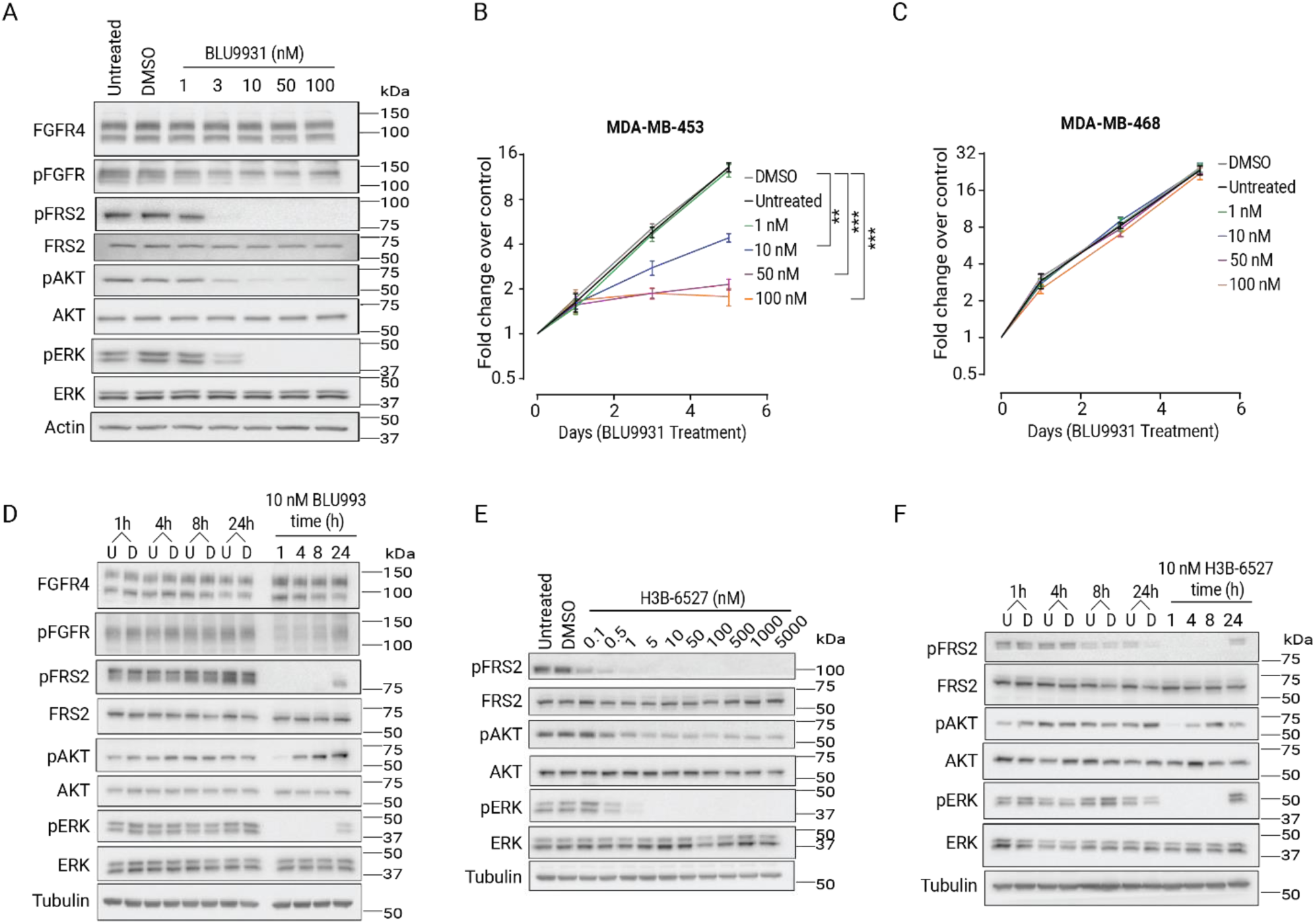
Effect of the FGFR4 inhibitor BLU9931 on FGFR4 downstream signalling pathways and proliferation in the MDA-MB-453 cell line. **(A)** Dose dependent effect of BLU9931 on expression and activation of downstream signalling proteins in MDA-MB-453 cells, 1 h post-treatment with the indicated doses. Effect of BLU9931 on proliferation in the FGFR4-positive MDA-MB-453 **(B)** and FGFR4-negative control MDA-MB-468 cell lines **(C)**. Cell proliferation was determined by direct cell counting. Error bars: mean ± standard error of three biological replicates. ** indicates p-value of < 0.01 and *** < 0.001, comparing individual BLU9931 concentrations to the DMSO vehicle control. U indicates untreated control, D indicates DMSO vehicle control. **(D)** Time course analysis of BLU9931 treatment. Expression and activation of downstream signalling proteins 1, 4, 8 and 24 h post-treatment with 10 nM of BLU9931. **(E)** Dose dependent effect of H3B-6527 on expression and activation of downstream signalling proteins in MDA-MB-453 cells, 1 h post-treatment with the indicated doses. **(F)** Time course analysis of H3B-6527 treatment. Expression and activation of downstream signalling proteins 1, 4, 8 and 24 h post-treatment with 10 nM of H3B-6527. Representative of three biological replicates. Note that the pFGFR signals were detected using a pan-phospho-FGFR antibody.

To understand how FGFR4 inhibition temporally influences downstream signalling, MDA-MB-453 cells were treated with 10nM of BLU9931 for 1, 4, 8 and 24 h (Fig. 1D). The levels of pFRS2 and pERK were initially decreased, followed by some signal recovery at 24 h (Fig. 1D). However, pAKT levels decreased after 1 h of drug treatment but the signal recovered at 4 h and then increased at 8 h and 24 h relative to the vehicle control (Fig. 1D). To identify if the bounce-back of pAKT is independent of the inhibitor used, MDA-MB-453 cells were treated with another FGFR4 inhibitor, H3B-6527, a specific FGFR4 inhibitor currently undergoing clinical trials for solid tumours (NCT02834780, NCT03424577). This resulted in the same pAKT rebound effect at the same time points (Fig. 1D-F). This indicates a general rewiring phenomenon following FGFR4 inhibition in MDA-MB-453 cells.

### A new mechanistic model of the integrated FGFR4 signalling network

To achieve a holistic and systematic understanding of drug-induced network rewiring, we developed a novel mechanistic mathematical model of the integrated FGFR signalling network. (Fig. 2A, S2). The model scope and assumptions along with supporting evidence are described in detail in Supplementary Text S1. The model was formulated using ordinary differential equations (ODEs), which describe the time-dependent rates of change of each species in the network based on the underlying biochemical reactions and their kinetics (see Materials and Methods). Detailed description of the reaction rates, kinetic laws used, and full set of ODEs are provided in Supplementary Tables S1-2.

**Figure 2.**
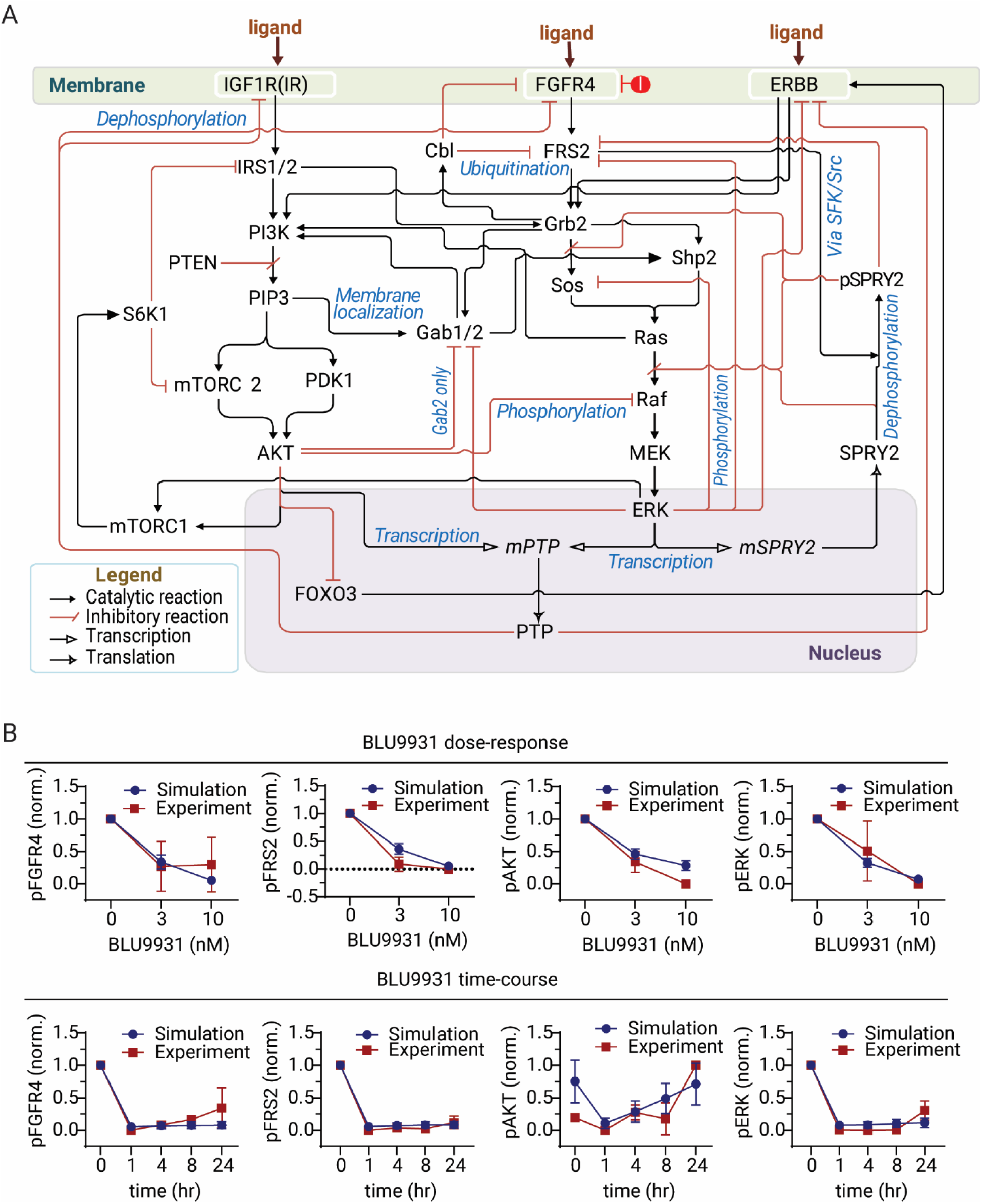
Development and calibration of a mathematical model of the FGFR4 signalling network. **(A)** A simplified schematic diagram depicting the network interactions included in the FGFR4 model, consisting of major RTKs, converging downstream signalling pathways and key feedback and crosstalk mechanisms. A detailed model reaction diagram is provided in Figure S2. The full model reaction rates and ODEs are given in Supplemental Table S1-2. **(B)** Model fitting to experimental data. Comparison of model simulation using best-fitted parameter sets (blue lines, error bars: mean ± standard error, n=50) against experimental time-course and dose-response data (red lines, error bars: mean ± standard error, n≥2) demonstrate good agreement between simulation and data. Data used for model calibration were quantified from Figures 1A, D.

To render our model context-specific and predictive, we first trained it using combined time- and dose-dependent signalling datasets from MDA-MB-453 cells, measured by Western blot (Fig. 1A,D). This was achieved by employing genetic algorithm-based optimisation procedures (Supplementary Text S1), which involved estimating the model parameters to ensure that model simulations fit the experimental data. Next, we conducted model identifiability analysis (21–24) to test if unique best-fitted parameter values could be obtained (see Materials & Methods and Supplementary Text S1). The analysis revealed partial identifiability (Fig. S3-4); i.e. at least one parameter remained non-uniquely identified, which aligns with the complexity of our model (24,25). This implies existence of multiple optimal parameter sets that fit the data similarly. We iterated the optimisation process 300 times using a genetic algorithm, yielding 50 equally well-fitted parameter sets (Supplementary Data S2-4). These sets were collectively used for simulations, which displayed strong concordance with the quantified experimental data (Fig. 2B). This confirms that the calibrated model, hereafter referred to as *model-1*, effectively reproduces the data both qualitatively and quantitatively.

### Concomitant inhibition of AKT, but not PI3K, eliminates pAKT rebound and AKT co-targeting enhances inhibition of MDA-MB-453 cell proliferation

To determine whether the rebound in AKT activation following FGFR4 inhibition is dosage-dependent, we utilised model-1 to simulate the temporal response of pAKT to escalating doses of BLU9931. Interestingly, the model predicted that high doses of BLU9931 induced an even stronger rebound of pAKT than low doses, with the effect being more pronounced at later time points (Fig. 3A). Given that PI3K is a direct upstream kinase of AKT signalling, we asked if co-targeting FGFR4 with a PI3K inhibitor could eliminate the rebound in pAKT. Model simulations showed that while combined FGFR and PI3K inhibition significantly suppressed pAKT levels, it did not completely eliminate the rebound pattern of pAKT (Fig. 3B).

**Figure 3.**
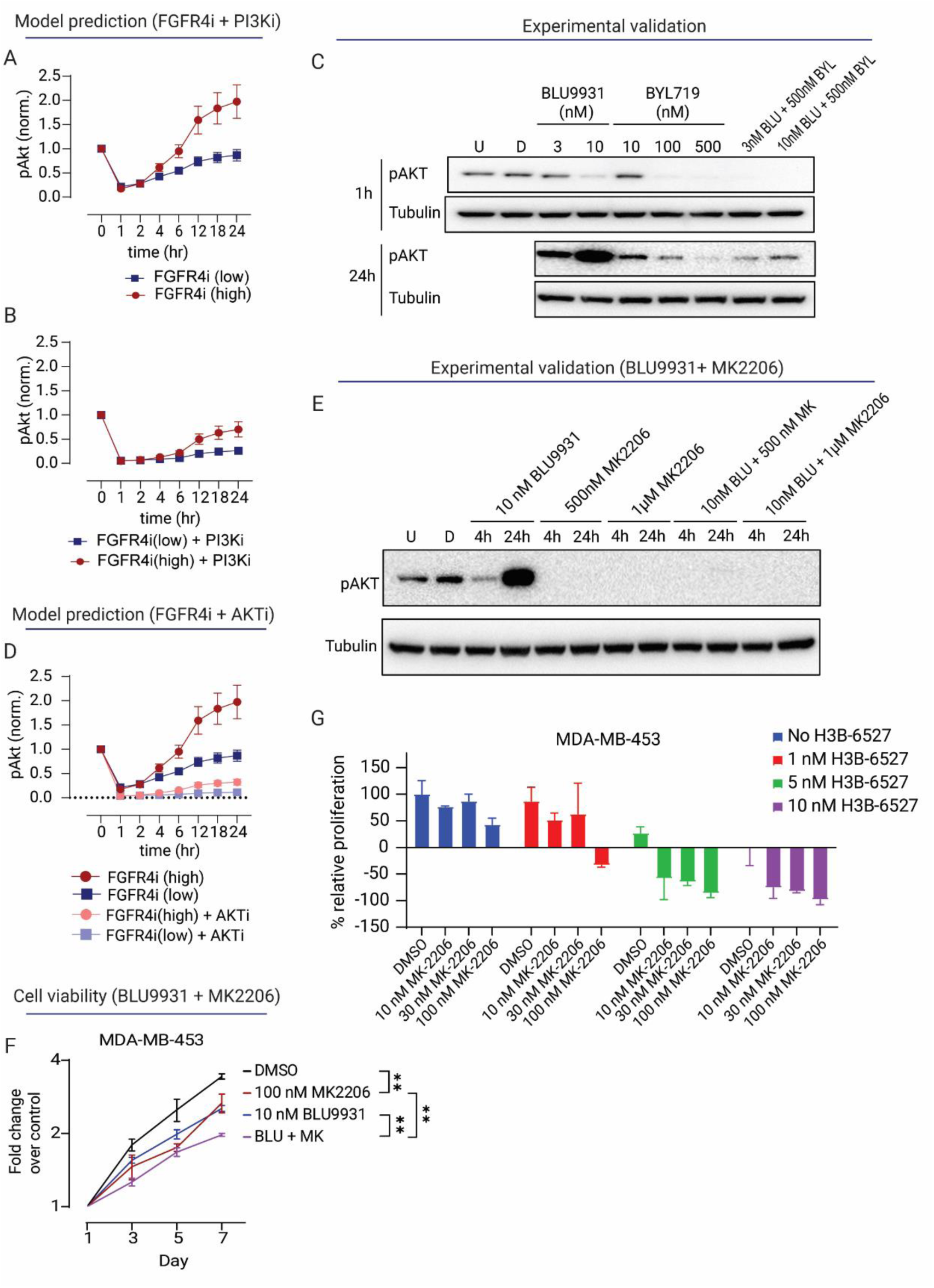
Co-inhibition of FGFR4 and AKT, but not PI3K, eliminates pAKT rebound and suppresses MDA-MB-453 cell proliferation. **(A-B)** Model prediction of phosphorylated AKT (pAKT) dynamics in response to FGFR4 inhibitor BLU9931 and PI3Kα inhibitor BYL719 in single or combination treatment. Error bars: mean ± standard error of 50 best-fitted parameter sets. We used the IC99 (FGFR4i (high)) and IC90 concentration (FGFR4i (low)) to inhibit FGFR4. For the inhibition of PI3K we used the IC90 concentration. **(C)** Dose dependent effect of BLU9931 in combination with BYL719 on AKT phosphorylation in MDA-MB-453 cells. pAKT was characterized 1 h and 24 h post-treatment with the indicated treatments. U indicates untreated control, D indicates DMSO vehicle control (represenative of 3 independent experiments). **(D)** Model prediction of pAKT dynamics in response to FGFR4 and AKT inhibitors in single or combination treatments. We used the IC99 (FGFR4i(high)) and IC90 concentration (FGFR4i(low)) to inhibit FGFR4. For the inhibition of AKT we used the IC90 concentration. Error bars: mean ± standard error of 50 best-fitted parameter sets. **(E)** Dose dependent effect of BLU9931 in combination with AKT inhibitor MK-2206 on pAKT in MDA-MB-453 cells. pAKT was characterized 1 h and 24 h post-treatment with the indicated treatments. U indicates untreated control, D indicates DMSO vehicle control (represenative of 3 independent experiments). **(F)** MDA-MB-453 cells were subjected to single inhibitor treatment or a combination of BLU9931 and MK-2206. Cell proliferation was determined by MTS assay. Error bars: mean ± standard error of four biological replicates. ** indicates p-value of < 0.01. **(G)** Combined drug effect of FGFR4 inhibitor H3B-6537 and AKT inhibitor MK-2206 on cell proliferation in the MDA-MB-453 cell line. Cell proliferation was determined by MTS assay. Error bars: mean ± standard error of three biological replicates.

To experimentally validate these model predictions, we first selected specific concentrations of the PI3Kα-specific inhibitor BYL719 to combine with BLU9931 (Fig. S5). Next, MDA-MB-453 cells were treated with BLU9931 and BYL719 alone or in combination for 1 and 24 h (Fig. 3C). The highest dose of BLU9931 and BYL719 decreased pAKT at 1 h, as did all the combination treatments (Fig. 3C). However, consistent with our model prediction, the higher dose of BLU9931 triggered a stronger rebound of pAKT at 24 h (Fig. 3C). Moreover, while the BLU9931 + BYL719 combination treatment suppressed the pAKT signal recovery, it did not completely eliminate the rebound, as predicted by our model.

Model simulations predicted that co-inhibition of AKT would strongly suppress the phosphorylation of AKT, as expected (Fig. 3D). To validate this experimentally, MDA-MB-453 cells were treated with BLU9931 and AKT inhibitor MK2206 alone or in combination for 4 h and 24 h (Fig. 3E, Fig. S6). The BLU9931 + MK2206 combination treatment potently suppressed pAKT at 24 hr, in agreement with our model prediction (Fig. 3D). MK2206 treatment alone also significantly suppressed pAKT to a greater extent than BLU9931 alone at 4 hr (Fig. 3E). In line with these findings, treatment with MK2206 significantly decreased the proliferation of MDA-MB-453 cells (Fig. 3F). Moreover, combining BLU9931 with MK2206 was significantly more effective in blocking cell proliferation than either agent alone (Fig. 3F). This work was further developed using the FGFR4 inhibitor H3B-6527. MDA-MB-453 cells were treated with different dosages of H3B-6527 and MK-2206 alone or in combination. Consistent with the BLU9931 data, the combined H3B-6527 + MK-2206 treatments caused synergistic effects on cell viability, confirming that the effect is independent of the FGFR4 inhibitor used (Fig. 3G). Drug synergism was quantified using the Bliss Independence and Loewe methods (Fig. S7A-B).

The prediction that inhibiting AKT is more effective than inhibiting PI3K in eliminating the pAKT rebound, despite the same target inhibition levels and PI3K being upstream of AKT, was intriguing (Fig. S8A). To gain further insight into this observation, we first performed model-based sensitivity analysis and identified network links that control the differential effects of PI3Ki versus AKTi on FGFR4i-induced pAKT reactivation. The sensitivity analysis (Fig. S8B) and time-course simulations (Fig. S9) demonstrated that the links related to negative feedback regulation, including the CBL-, SPRY2-, and PTP-mediated negative feedback loops, enhanced the differential effect of AKTi and PI3Ki on pAKT rebound (red line, Fig S8C). Conversely, the links associated with GAB proteins and positive feedback reduced the differential effects (blue line, Fig S8C). Next, we analysed the correlation between the magnitude of pAKT rebound and the differential effects of AKTi and PI3Ki. For this, we introduced the rebound (RB) score to describe the early response of pAKT to FGFR4i, followed by a rebound at a later time (Fig. S8D). We found that a stronger pAKT rebound exaggerates the differential effect between AKTi and PI3Ki on suppression of pAKT, as shown in Fig. S8E. This indicates that when the FGFR4-induced pAKT rebound is more pronounced, AKT is a more suitable co-target of FGFR4 than PI3K.

Collectively, our integrative studies indicated that the efficacy of FGFR4 inhibitors in this TNBC cell line is limited by the rebound of AKT activity, which can be more effectively inhibited by co-targeting AKT rather than PI3K.

### The role of specific RTKs in modulating FGFR4i sensitivity

To further investigate the dynamic signalling remodelling following FGFR4 inhibition, we enhanced our mathematical model by including additional time-course and dose-response signalling data from combined FGFR and PI3K inhibition experiments (Fig. S10). These datasets, previously excluded from model-1, helped recalibrate the model. The refined model-2 accurately replicated all training data using a new suite of 50 best-fitted parameter sets (Fig. S11; model identifiability analysis results in Figs S12-13).

Next, we extended our model simulation readouts beyond pAKT to include phosphorylated IRS, IGF1R and ErbBs in response to FGFR4i, as these proteins are regulated by multiple negative feedback loops (Fig. 2A). Model simulations predicted strong upregulation of pIGF1R, pIRS and pErbB at 24 hr following BLU9931 treatment at both low and high doses (Fig. 4A). Interestingly, higher BLU9931 concentrations triggered stronger upregulation of these proteins (Fig S14), as seen previously for pAKT (Figs. 3A-B). These predictions suggest that FGFR4 inhibition remodelled the RTK network in TNBC cells, causing pAKT rebound and compromising the inhibitory effect of FGFR4i on cell proliferation. To test this hypothesis and the model predictions, we assessed the expression and phosphorylation of specific RTKs in MDA-MB-453 cells at 1 hr and 24 hr after BLU9931 treatment. Consistent with the model predictions, we observed significant upregulation of pIGF1R, pIRS, and pErbB2-3 at 24 hr compared to the control or 1 hr treatment (Fig. 4B).

**Figure 4.**
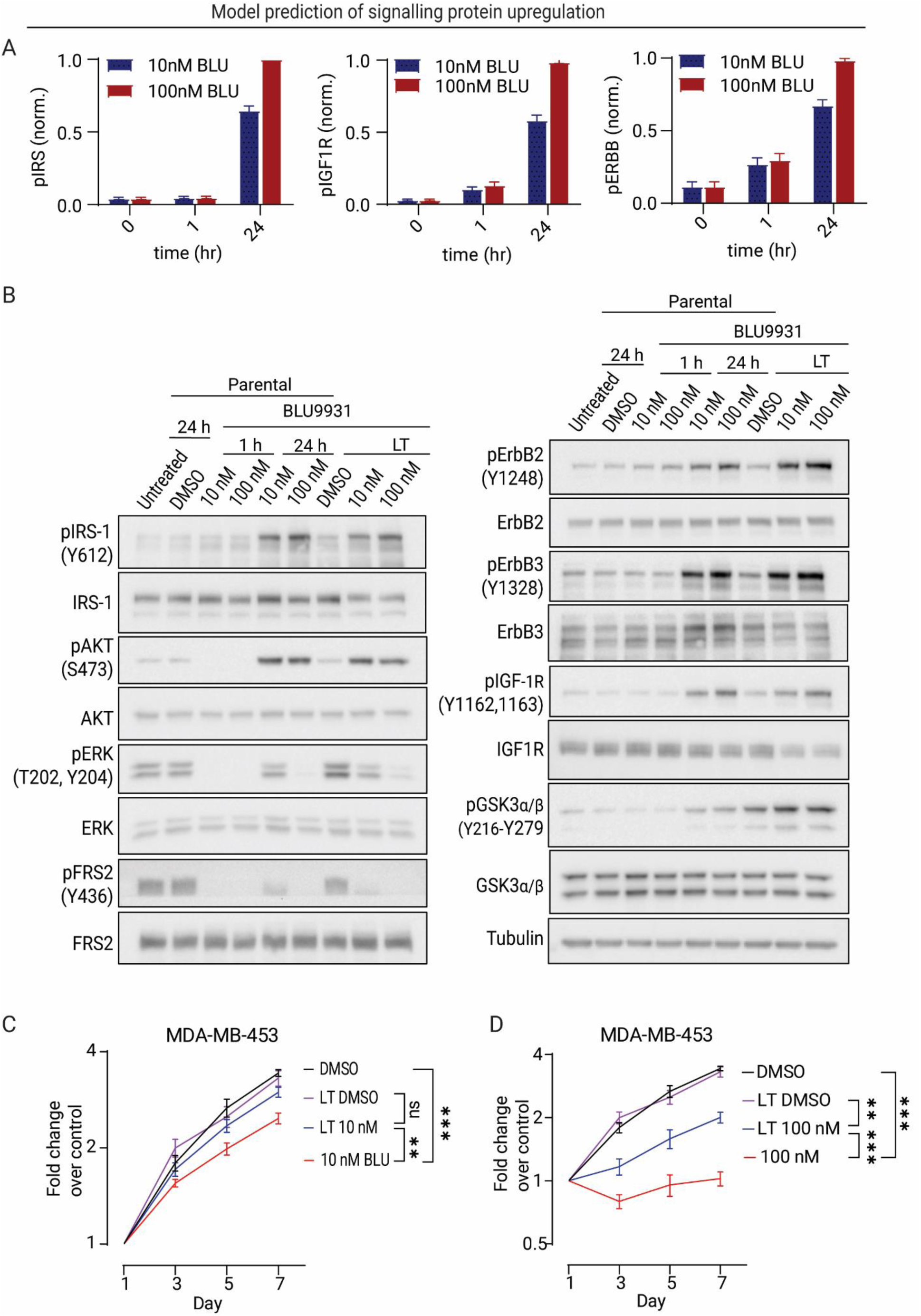
Identification of upregulated RTKs in parental and BLU9931-resistant MDA-MB-453 cells. **(A)** Model prediction of the dynamic responses of specific signalling proteins, predicting they are temporally upregulated following BLU9931 treatment. Error bars: mean ± standard error of 50 best-fitted parameter sets. **(B)** Expression and activation of RTKs and FGFR4 downstream signalling proteins at 1 and 24 h post-treatment with the indicated BLU9931 concentrations in parental MDA-MB-453 cells, compared with long-term (LT) BLU9931-resistant MDA-MB-453 cells. The latter were harvested 24 h post-seeding. **(C-D)** The proliferation of parental and long-term BLU9931-resistant MDA-MB-453 cells treated with DMSO vehicle control, 10 nM (C) or 100 nM BLU9931 (D). Cell proliferation was determined by MTS assay. Error bars: mean ± standard error of six biological replicates. ** indicates p-value of < 0.01, *** < 0.001.

In addition to investigating the short-term remodelling of the FGFR4 network in response to BLU9931, we also interrogated its long-term effects. To this end, we exposed MDA-MB-453 cells to BLU9931 for at least 3 months to allow isolation of resistant cells (see Material and Methods). Long-term drug-treated cells showed no significant growth inhibition at 10 nM BLU9931 compared to parental cells (Fig. 4C), while 100 nM of the drug reduced growth, although this potency was stronger in parental cells (Fig. 4D). To investigate the mechanism underlying long-term resistance to BLU9931, we sequenced the FGFR4 kinase domain and characterized the expression profile of FGFR4 signalling network components in parental and long-term BLU9931-treated cells. FGFR4 gatekeeper mutations (26) were not detected (Fig S15), and similar to parental cells treated for 24 h, long-term BLU9931-treated cells showed a marked upregulation of pAKT, pIGF1R, pIRS, and pErbB2-3 (Fig. 4B). Furthermore, levels of phosphorylated GSK3α/β and pY1248 ErbB2 were higher in long-term BLU9931-treated cells than in parental cells treated for 24 hrs. While increased activation of particular RTKs induced by 24-hour BLU9931 treatment is maintained in the long-term drug-refractory cells, additional changes likely underpin long-term resistance to BLU9931. These data suggest potential targets in addition to AKT for combination therapies with FGFR4 inhibition, which will be addressed next.

### Model prediction and validation of synergistic drug combinations co-targeting FGFR4

Our computational model provides a powerful framework for the systematic prediction of nodes that could be co-targeted with FGFR4 to overcome adaptive resistance to anti-FGFR4 monotherapy. To achieve this, we further refined model-2 by incorporating the new experimental data from Fig. 4B and S16 for model recalibration, evolving it to become *model-3*. The refined model reproduces all the training data well (Fig. S17), and displays improved identifiability compared to the previous models (Figs. S18-20). Using model-3, we conducted in silico simulations to evaluate the efficacy of 19 possible pair-wise drug combinations co-targeting FGFR4 and 19 other network nodes (Fig. 5A). Highest Single Agent (HSA) and the coefficient of drug interaction (CDI) were computed to evaluate the theoretical performance of each combination in suppressing cell proliferation relative to single-drug treatments (Fig. 5A and S21A, also see Materials and Methods). By sorting these synergy scores, we were able to rank and prioritise the drug combinations based on their predicted synergistic effects (Fig. 5A and S21A).

**Figure 5.**
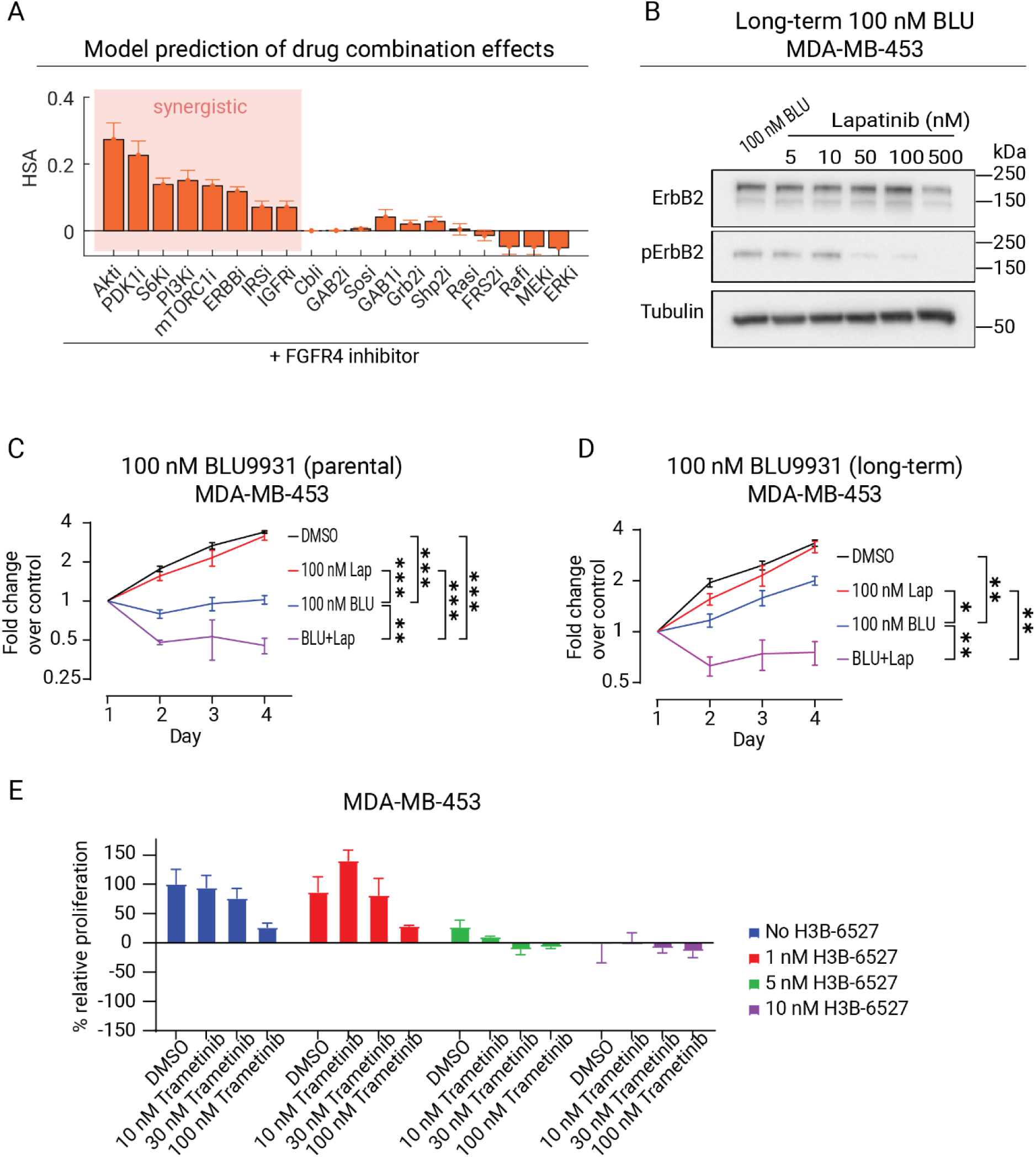
Model prediction of synergistic drug combinations and experimental validation. **(A)** In silico prediction of the effect of 19 possible pair-wise drug combinations co-targeting FGFR4 and various network components, assessed using drug synergy scores of HSA. The value that is >1, =1, <1 indicate synergistic, additive or antagonistic effects, respectively; and higher HSA scores indicate stronger synergism (Materials and Methods). Error bars: mean ± standard error of 50 best-fitted parameter sets. **(B)** Dose dependent effect of Lapatinib on the expression and phosphorylation of ErbB2 in long-term 100 nM BLU9931-resistant MDA-MB-453 cells. **(C-D)** Proliferation of parental and long-term 100 nM BLU9931-resistant MDA-MB-453 cells subjected to single inhibitor or combination treatment of BLU9931 (BLU) and Lapatinib (Lap) at the indicated doses. Cell proliferation was determined by MTS assay. **(E)** Combined drug effect of FGFR4 inhibitor H3B-6537 and MEK inhibitor trametinib on cell proliferation of the MDA-MB-453 cell line. Cell proliferation was determined by MTS assay. Error bars: mean ± standard error of three biological replicates. * indicates p-value of <0.05, ** < 0.01, *** < 0.001.

Our model predicted synergistic drug combinations for co-targeting FGFR4, with dual FGFR4 and AKT inhibition showing the most synergistic effect (Fig. 5A). In contrast, co-targeting FGFR4 and MEK was predicted to be non-synergistic. Interestingly, co-targeting ErbB receptors was predicted to be synergistic (Fig. 5A). Our data demonstrating upregulation of ErbBs and IGF1R in response to FGFR4 inhibition (Fig. 4) are consistent with our model’s prediction that ErbBs would be a synergistic FGFR4 co-target despite being in a different signalling branch than the IGF1R-PI3K-AKT cascade. To validate this prediction, we treated long-term BLU9931-maintained cells with Lapatinib, which predominantly targets ErbB2 in MDA-MB-453 cells since EGFR/ErbB1 expression is low (27), and selected 100 nM as the most appropriate dose for subsequent experiments (Fig. 5B). Lapatinib treatment alone had no effect on parental or resistant cells (Fig. 5C-D) but the combination treatment of BLU9931 and Lapatinib induced cell death in both parental and resistant cells (Fig. 5C-D). Note that at 10 nM BLU9931, the combination treatment with Lapatinib restored growth inhibition only in cells maintained long-term (Fig. S21B-D). To further confirm the model prediction of non-synergistic combination, we conducted an experiment where MDA-MB-453 cells were treated with different doses of H3B-6527 and MEK inhibitor trametinib, either alone or in combination, and their effects on cell viability were assessed (Fig 5E). Interestingly, unlike the synergistic effects observed with the combined H3B-6527+MK-2206 treatments (Fig. 3G), the combination of H3B-6527+trametinib failed to exhibit any synergistic effects on cell viability inhibition for almost all dose combinations (Fig. 5E and S22).

Together, these results indicate that long-term BLU9931-maintained cells are more dependent on ErbB2, consistent with the enhanced ErbB2 Y1248 phosphorylation (Fig 4B), and demonstrate that co-targeting ErbB2 synergistically enhances the efficacy of FGFR4 inhibition in MDA-MB-453 TNBC cells.

### Analysis of network connections that drive pAKT and pErbB rebounds

Our analyses identified RTKs such as ErbBs and IGF1R as potential mediators of pAKT rebound following FGFR4i treatment. However, the network features that drive this rebound remain unclear. To address this issue, we conducted a sensitivity analysis using model-3 where we systematically blocked network links and observed the resulting changes in pAKT rebound following BLU9931 treatment (Fig. 6A). While a visual assessment indicated strong and varied changes to the pAKT dynamics depending on the perturbed model parameters (Fig. 6A), to assess these changes more accurately we computed the rebound metric introduced earlier (*RB*, Fig. S8D) for each of the perturbed and control conditions (Fig. 6B).

**Figure 6.**
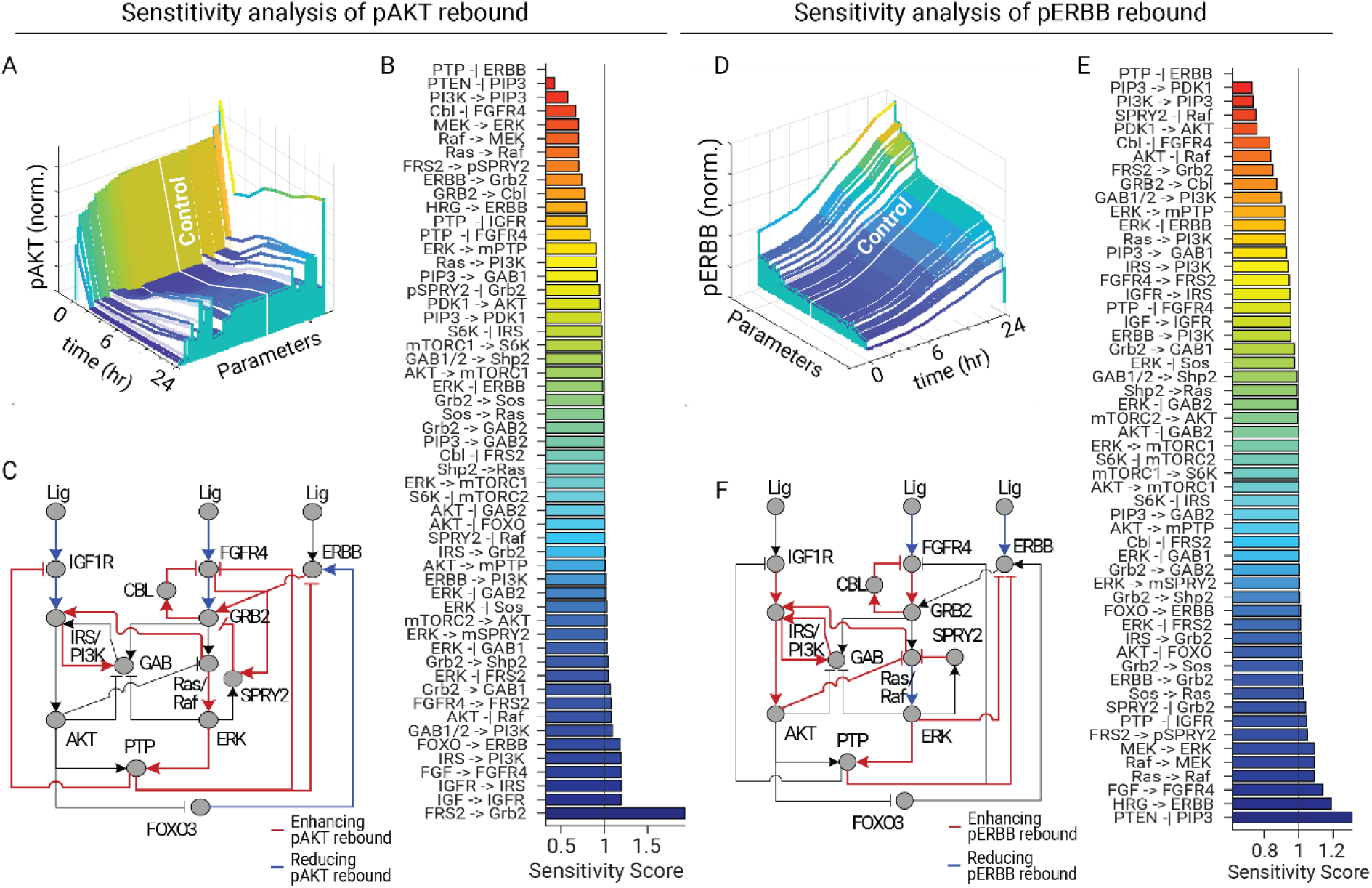
Model-based sensitivity analysis and identification of network architecture controlling pAKT and pErbB rebound. **(A, D)** A 3D plot (waterfall plot) showing in silico simulation of the temporal phosphorylated AKT and ErbB dynamics in response to FGFR4 inhibition. The dynamics of the intact (non-perturbed) model is highlighted in white as a control. The pAKT and pErbB time profiles were averaged across 50 best-fitted parameter sets. **(B, E)** Changes in the RB score of BLU9931-induced pAKT and pErbB response were measured by a sensitivity score, relative to the control (no perturbation) condition. If the sensitivity score is less than 1, it indicates that the parameters promote the reactivation of pAKT and ErbB. If the score is greater than 1, it indicates that the parameters inhibit it. **(C, F)** Simplified network models and key mechanisms that play a critical role in inducing the reactivation of pAKT and pErbB.

We found that inhibiting certain parameters decreased the RB score, indicating a positive role for the corresponding network links in mediating pAKT rebound (Fig. 6B). In contrast, for almost a quarter of the remaining parameters, blocking them had the opposite effect, while the remaining three-quarters did not have a detectable impact on the RB score (Fig. 6B). By mapping these parameters onto the network interaction diagram, we found that the magnitude of the pAKT RB score is under the positive control of the negative feedback loops mediated by CBL and PTP, with the PTP-to-ErbB feedback loop having the most significant impact (Fig. 6B-C). As these feedback loops act primarily upon the RTK receptors, including ErbB and IGF1R, the findings align with our prior experimental results showing the upregulation of these RTKs following FGFR4 inhibition (Fig. 4). On the other hand, we found that network links that had a negative impact on the pAKT RB score tended to be positioned along the core RTK signalling axes, such as ligand-to-IGFR1/FGFR4, which promote the activation of the RTKs and downstream signals (Fig. 6C).

To further investigate the mechanism of ErbB signal reactivation since this is likely associated with the pAKT rebound, we conducted a similar sensitivity analysis on the dynamics of pErbB. The result revealed that, similar to pAKT rebound, the links related to PTP- and CBL-mediated negative feedback positively regulated the ErbB RB score (Fig. 6D-F). However, we observed that the ERK-to-ErbB negative feedback had an impact only on ErbB reactivation, indicating that these links may be more specific to the ErbB signalling pathway (Fig. 6F). Together, these systematic analyses identify key network features that drive AKT and ErbB signal reactivation, providing important insights into the mechanism underlying the signalling dynamic remodelling following FGFR4 inhibition.

### Diverse pAKT/pERK response profiles following FGFR4 inhibition across cancer cell lines

In our previous work (28), we showed that cell-to-cell variability in protein expression levels is a major source of heterogeneity in signalling response. To explore the hypothesis that FGFR4i-induced signalling dynamics are dictated by the protein expression profiles across specific cancer cell types, we obtained protein expression data from the Cancer Cell Line Encyclopedia (CCLE) consortium for 350 different cancer cell lines representing diverse tissue origins (29). We utilised this data to customize our MDA-MB-453 based model-3, generating 350 models that reflect the specific protein expression profile of each cell type (see Fig. S23A for detailed workflow). By simulating the temporal response of pAKT and pERK after FGFR4 inhibition using these cell type-specific models, we identified a remarkable heterogeneity in the response of these outputs, ranging from no significant change to strong rebound to straight increase patterns for pAKT (Fig. 7A), while pERK predominantly displayed rebound or monotonic decrease patterns (Fig. 7B).

**Figure 7.**
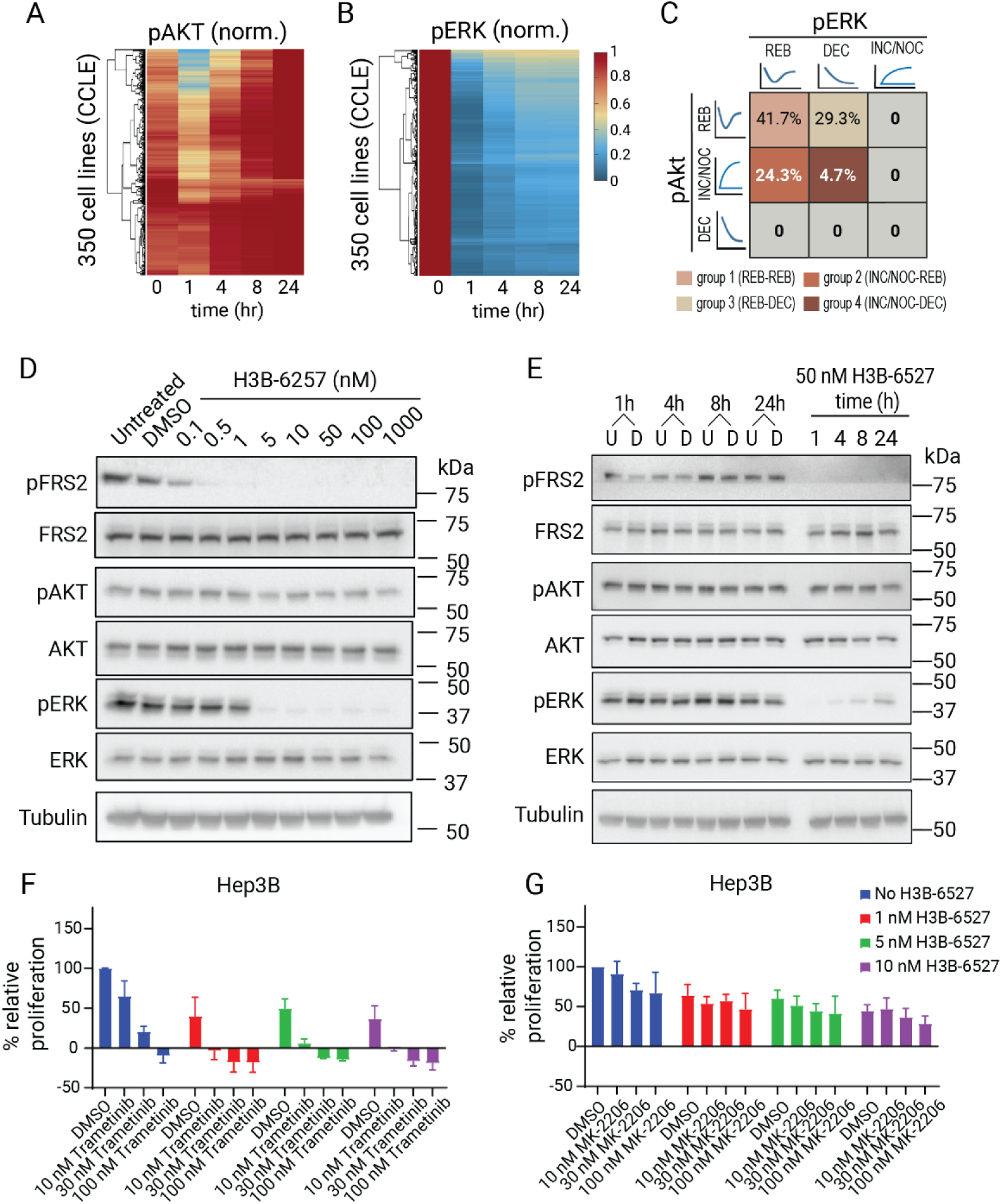
Analysis of drug-induced pAKT/pERK response profiles across a large panel of cancer cell types. **(A, B)** Model prediction of pAKT and pERK time-course profiles in response to FGFR4 inhibition in 350 cancer cell lines using cell-type specific models generated by incorporating protein expression data from CCLE. **(C)** Classification of the cell lines into subgroups based on the response patterns of pAKT and pERK. INC: increase, DEC: decrease, REB: rebound, NOC: no change. **(D)** Dose dependent effect of H3B-6527 on expression and activation of FGFR4 downstream signalling proteins in Hep3B cells 1 h post-treatment with the indicated doses. **(E)** Time course analysis of treatment using the FGFR4 inhibitor H3B-6527 on FGFR4 downstream signalling pathways in the Hep3B cell line. Expression and activation of downstream signalling proteins were characterized 1, 4, 8 and 24 h post-treatment with 50 nM H3B-6527. U indicates untreated control, D indicates DMSO vehicle control. Representative of three biological replicates. Combined drug effect of FGFR4 inhibitor H3B-6537 and MEK inhibitor Trametinib (**F**) or AKT inhibitor MK-2206 (**G**) on cell proliferation in the Hep3B cell line. Error bars: mean ± standard error of three biological replicates. Cell proliferation was determined by MTS assay.

We defined three composite patterns (REB: rebound; INC/NOC: increase or no significant change; and DEC: decrease) and used them to classify the cell lines into four major subgroups (namely groups 1-4: REB-REB; INC/NOC-REB; REB-DEC; and INC/NOC-DEC) based on their distinct response profiles of pAKT/pERK respectively (Fig. 7C and Fig. S23B). Our simulations revealed that FGFR4 inhibition induced pERK rebound in about two-thirds of the cell lines (66%), indicating that reactivation of ERK represents a salient response feature post FGFR4i treatment in a subset of cell types that likely blunts the treatment effect. Although pAKT displayed rebound in about 71% of the cell lines, our simulations suggest that FGFR4 inhibition either did not significantly affect or induced a straight increase of pAKT in the remaining cell lines (29%). Notably, only 5% of the cell lines exhibited concomitant pAKT increase/no change and pERK decrease, while 42% exhibited rebound in both pAKT and pERK (Fig. 7C).

Next, to examine whether the modulation of network components could potentially trigger a transition between different response groups, we employed the MDA-MB-453 model (group 3) and systematically altered the expression levels of each component. The perturbation analysis indicated that downregulation of FGFR4, FRS2, Grb2, or Ras, or upregulation of PDK1, AKT, PIP2 or PI3K transitioned pAKT from a rebound state to an increase or no change state, resulting in a group 3 → 4 transition (Fig. S24).

Together, these studies highlight the heterogeneity and adaptability of signalling responses to FGFR4-targeted therapy in cancer cells. These findings underscore the necessity of considering the network context when analysing drug-induced network remodelling and carry implications for the development of personalized cancer therapies.

### Alternative network rewiring following selective FGFR4 inhibition in Hep3B cells

Our model-based analysis predicted distinct cancer cell subgroups with different response patterns for pAKT-pERK, which can potentially limit the effect of FGFR4 inhibition. The MDA-MB-453 cell line represents group 3 (Fig. 1D, F), and to validate the model prediction of group 2, we identified Hep3B, an HCC cell line with active FGFR4 signalling due to FGF19 gene amplification (9), as a member of this group.

Western blot analysis of Hep3B cells treated with H3B-6527 revealed a significant decrease in phosphorylated FRS2 and ERK after 1 hr (Fig. 7D, E). However, we also observed an early and temporally increasing rebound of pERK levels, despite the durable suppression of pFRS2 (Fig. 7 E). Notably, pAKT showed no significant change after H3B-6527 treatment in these cells (Fig. 7E and Fig. S25). These dynamics are consistent with the predicted behaviour of group 2, confirming the validity our model-based approach for identifying cell type-specific response patterns to anti-FGFR4 treatment. For the Hep3B model, downregulation of ERK, MEK, Raf, or Ras, or upregulation of AKT transitioned pERK from a rebound state to a decrease, triggering a group 2 → 4 transition (Fig. S26).

The H3B-6527 treatment-induced rebound of pERK suggests that co-targeting ERK may enhance the sensitivity of Hep3B cells to H3B-6527. To test this hypothesis, we first determined the appropriate concentration of trametinib, an inhibitor that targets ERK’s direct upstream kinase MEK, to be used in combination treatments. Treatment with 5nM or higher of trametinib led to decreased pERK in Hep3B cells (Fig. S27). We then tested the effects of combined H3B-6527 and trametinib in comparison to single-drug treatments. While H3B-6527 and trametinib alone decreased Hep3B cell proliferation (Fig. 7F), their combination resulted in strong synergistic effects (Fig. 7F, S28A-B). Interestingly, when we treated the cells with H3B-6527 and the AKT inhibitor MK-2206, we did not observe strong synergistic effects between them (Fig. 7G and S28C-D). This observation is consistent with the fact that pAKT was not significantly affected by H3B-6527 treatment (Fig. 7G). These results indicate that the efficacy of H3B-6527 is improved by blocking the ERK signal bounce-back and highlights the combination of H3B-6527 and trametinib as a promising therapeutic approach for HCC with this network behaviour.

## Discussion

In this study we develop a comprehensive computational model for FGFR4 signalling and shed new light on the mechanisms utilised by cancer cells to counter FGFR4 inhibition. Furthermore, we identify heterogeneity in tumour cell responses to FGFR4 inhibition and based on this, highlight potential context-specific combined drug strategies for enhanced personalised treatment outcomes.

Our new computational model of the FGFR4 signalling network integrates key signalling cascades downstream of FGFR4 and is rigorously calibrated using diverse time-course and dose-response data sets, employing ordinary differential equation formalism. This represents a significant step forward from early models, which either failed to capture intracellular signalling (30–34) or neglected key downstream pathways (35,36). Some recent studies have adopted logic-based Boolean modelling (37); however, these cannot capture the complex temporal aspects of drug-induced signalling dynamics. Therefore, our model fills these critical gaps by providing a more detailed and comprehensive platform for the analysis and prediction of these dynamics.

The integration of our computational modelling and experimental analyses has shed light on the network rewiring occurring in breast cancer cells to resist FGFR4 inhibition. In MDA-MB-453 TNBC cells we determined that selective inhibition of FGFR4 leads to significant reactivation of the pro-survival protein AKT. However, an interesting predicted and validated phenomenon was that targeting AKT more effectively eliminated AKT rebound than inhibiting PI3K. Sensitivity analysis showed that blocking negative feedback mechanisms, such as CBL-, SPRY2-, and PTP-mediated loops, mitigated the differential effect of AKT and PI3K inhibitors on pAKT rebound. Further analysis revealed a correlation between the strength of pAKT rebound and the differential impact of AKT and PI3K inhibitors. Stronger pAKT rebound widens this differential effect, while its absence narrows it, indicating a dominant role for feedback loops like CBL and PTP in controlling the contrasting impacts of AKT and PI3K inhibitors on pAKT response. Although experimental validation of these predictions is pending, our computational findings have illuminated otherwise non-intuitive observations.

The focus of most research has been on combining FGFR4 inhibition with radio or chemotherapeutics, leaving the area of combinatorial targeting of FGFR4 relatively understudied (38,39). To expedite the identification of novel combination treatments that target FGFR4 we predicted the impact of pair-wise drug combinations in a systematic and unbiased manner. This highlighted ErbB receptors as FGFR4 co-targets in MDA-MB-453 cells, consistent with the activation of alternative RTKs as an escape mechanism previously reported in FGFR-resistance models (40,41). For example, FGFR3-dependent bladder cancer cell lines developed resistance to the pan-FGFR inhibitor BGJ398 by switching receptor signalling to ErbB2 or ErbB3 (40).

Given these insights, it is important to note that the mechanisms underlying adaptive resistance can vary significantly depending on the context, even when the same drug is used (42). This variability is driven by differences in protein expression and mutational profiles among different cancer cell types, which influence feedback strength and crosstalk, ultimately dictating drug response (18,28). To address this, we incorporated cell type-specific protein expression data to generate hundreds of cell type-specific computational models, which led to the identification of distinct response groups that exhibit unique behaviours after FGFR4 inhibition. We identified and experimentally validated the HCC cell line Hep3B as a cell type exhibiting pERK rebound with no significant change in pAKT. Importantly, co-targeting FGFR4 and MEK, upstream of ERK, in Hep3B cells significantly and synergistically enhanced the inhibition of cell proliferation, highlighting this approach as a promising precision treatment strategy for HCC. The unique clustering of response dynamics uncovered here underscore the need to consider network context in drug resistance analysis, and has potential implications in clinical settings, especially in the context of personalized cancer therapies.

We also demonstrated that manipulation of network interference points could shift response behaviours, such as transitioning from rebound to progressive increase in pAKT activation. This illustrates how cancer cells can exploit the plasticity of signalling networks to evolve from drug-sensitive to drug-resistant states. Furthermore, this new knowledge can assist in designing new therapeutic strategies to counteract drug-induced reactivation of pro-growth signalling by transforming pAKT/pERK (re)activation into a reduction in activity. These findings highlight the effectiveness of using computational modelling in identifying nodal alterations that manipulate dynamic signalling behaviour. Adding to the novelty of our study, we introduced innovative metrics that enable systematic and quantitative detection and characterization of drug-induced adaptive rebound in signalling proteins, including the rebound (RB) metric. Not only does this offer a more precise definition of adaptive signalling rebound, but it also enables systematic comparison across various conditions. Indeed, the utilization of these metrics in our study has facilitated model sensitivity analysis of potential mechanisms underlying the rebound of AKT and ErbB activities following FGFR4 inhibition, which would have been challenging otherwise.

In conclusion, by taking an integrative approach, we have uncovered the mechanisms that drive FGFR4 network rewiring and discovered potential combinatorial targeted therapies that deliver improved efficacy for FGFR4-driven TNBC and HCC cancers. Our finding that co-targeting FGFR4 and ErbB2 was particularly effective suggests that this targeted treatment strategy could extend to other breast cancer subtypes with poor prognosis, including luminal B and HER2 breast cancers. Furthermore, the computational models and integrative systems-based approach utilised in this study lay a robust foundation for future investigations into network rewiring and adaptive resistance related to targeting other RTKs.

## Materials and Methods

### Cell lines, cell culture and reagents

The TNBC cell line, MDA-MB-453 and HCC cell line, Hep3B were purchased from ATCC. MDA-MB-453 was cultured in RPMI-1640 (Gibco) supplemented with 10% (v/v) FBS (Moregate), 10 μg/mL Actrapid penfill insulin (Clifford Hallam Healthcare) and 20 mM HEPES (Gibco). Hep3B was cultured in EMEM (USbio) supplemented with 10% (v/v) FBS and 1 mM sodium pyruvate (Gibco).

### Inhibitors and treatment

The following inhibitors were purchased from Selleckchem: FGFR4 inhibitor BLU9931 and H3B-6527, ErbB family inhibitor Lapatinib, PI3Kα inhibitor BYL719, AKT inhibitor MK2206 and MEK inhibitor Trametinib. All inhibitors were reconstituted in DMSO.

For inhibitor treatment, cells were seeded into culture plates with an 80% end point confluence for all cell lines. After 24 h, cells were treated for the indicated times with the specific inhibitor or DMSO as vehicle control.

### Generation of FGFR4 inhibitor resistant cells

MDA-MB-453 cells were seeded into 10 cm plates at a density of 500 cells. After 48 h, cells were treated with DMSO (as vehicle control) or FGFR4 inhibitor BLU9931 for at least 3 months. Culture medium was replaced twice a week. When cells formed colonies visible to the naked eye, small pieces of sterile filter paper soaked in trypsin were used to detach cells which were collected for further maintenance in media containing DMSO or FGFR4 inhibitor. These cells were termed long-term BLU9931 MDA-MB-453 cells in this study.

### Immunoblotting

Protein lysates were prepared in RIPA buffer and subjected to Western blotting as described previously (43).

The following antibodies were purchased from Cell Signaling Technology: pan-phosFGFR (Y653/654) (3471), AKT (4685), ERK (4695), pAKT (S473) (4058), pERK (T202, Y204) (4370), pFRS2 (Y436) (3861), ErbB2 (2165), pErbB2 (Y1248) (2247), ErbB3 (4754), pErbB3 (Y1328) (8017), IGF-1R (9750) and GSK-3α/β (5676). The following antibodies were purchased from Santa Cruz Biotechnology: FGFR4 (sc-136988) and β-actin (sc-69879). The α-tubulin (T5168) and FRS2 (05-502) antibodies were purchased from Sigma-Aldrich. The pIRS-1 (Y612) (44-816G) and pIGF-1R (Y1162/1163) (44-804G) were purchased from Biosource. An IRS-1 (6248) antibody was purchased from Upstate and a pGSK-3α/β (Y216/279) (ab4797-50) was purchased from Abcam.

### Cell viability assays

Trypsinized cells were stained with Trypan blue (EVS-1000, NanoTek), then transferred to an EVE cell counting slide (EVS-1000, NanoTek) and counted with the EVE automatic cell counter (EVE-MC-DEMO, NanoTek) according to the manufacturer’s protocol. MTS proliferation assays were undertaken as previously described (43).

### RNA isolation, RT-PCR and Sanger sequencing

Total RNA was isolated from parental MDA-MB-453 and long-term BLU9931 MDA-MB-453 cells using a RNeasy mini kit (Qiagen) following the manufacturer’s protocol. RNA was quantified using a Nanodrop ND-1000 (NanoDrop Technologies). RNAs were reverse transcribed using a high-capacity cDNA reverse transcription kit (Thermoscientific). Subsequently, cDNA was amplified by PCR to identify gatekeeper mutations in the FGFR4 kinase domain using forward (F) primers and reverse (R) primers (Table S3). The PCR products were resolved by gel electrophoresis, and the bands at the predicted product size were excised and purified with a gel and PCR clean-up system (Promega). Sanger sequencing was completed by the Micromon facility at Monash University. Reactions were repeated on three biological replicates.

### Statistical analysis

Quantification of western blots by densitometry was performed using ImageLab version 5.2.1 (Bio-Rad) and statistical t-tests were performed using GraphPad Prism 8 and Microsoft-Excel.

### Computational modelling

The FGFR4-centered model was formulated using ordinary differential equations (ODEs). The rate equations and full set of ODEs are given in Supplementary Tables S1-2. The model construction and calibration processes were implemented in MATLAB (The MathWorks. Inc. 2023a) and the IQM toolbox (http://www.intiquan.com/intiquan-tools/) was used to compile the IQM file for a MEX file which makes the simulation faster. To facilitate model exchange, an exchangeable Systems Biology Markup Language (SBML) file of the model is provided as Supplementary Data S1-3. The code for the modelling has been deposited to Github and can be accessed at https://github.com/NguyenLabNetworkModeling/FGFR4-Signaling-Network-Model.git.

### Model calibration

The adequacy and specificity of a mathematical model is generally justified by its ability to recapitulate experimental data. This is achieved through model calibration (also referred to as model training or fitting) that involves estimation of the unmeasured model parameters so that simulations using these parameter values (called best-fitted parameter sets) could recapitulate the training data (Fig. S30-32). Here, parameter estimation was done by minimizing an objective function 𝐽 that quantifies the discrepancies between model simulations and the corresponding experimental measurements. A genetic algorithm was used to optimize the objective function (44–46), utilising the Global Optimization Toolbox and the function *ga* in MATLAB. Model calibration was carried out on a virtual machine consisting of 32 Intel Xeon 2.2 GHz processors running in parallel. A more detailed description of the model calibration process is given in Supplementary Text S1. As a result, we obtained 50 best-fitted parameter sets that were collectively used for model simulations (Fig. S32); these are given in Supplementary Data S2-4.

### Model-based simulation and computation of drug synergy

Because the synergy between two drugs can depend on the specific doses at which they are combined, we used IC50 value of individual drugs as a reference concentration for the simulation (see Supplementary Information for details). Using the simulated dose-response matrix, we evaluated in silico the efficacy and possible synergism of 19 possible combinatorial strategies co-targeting FGFR4 and each of 19 network components. Drug synergy was computed based on the coefficient of drug interaction (CDI) and highest single agent (HSA) metric (47–49): CDI = *R_12_*/(*R_1_*×*R_2_*), where *R_12_* is a normalized biological response (e.g., cell viability) by the combined treatment of drug 1 and 2, and *R_1_* and *R_2_* are the response by the single drug treatment, respectively. Note *R* = 1 (and 0) indicates no drug response (complete inhibition, respectively). CDI <1, = 1 or >1 indicates that the drugs are synergistic, additive or antagonistic, respectively. According to the HSA model, the synergistic effect is equal to the greater effect of individual drugs (49). Thus, HAS = *E_12_* – max(*E_1_*,*E_2_*). *E_12_* is a normalized drug effect by the combined treatment of drug 1 and 2, and *E_1_* and *E_2_* are the response by the single drug treatment, respectively. Note *E* = 0 (and 1) indicates no drug effect (maximum effect, respectively). For the readability and easy comparison between CDI and HAS, we used a log2-transformed CDI metric (-log2(CDI)): Thus, for the both metrics the score > 0, = 0 or < 0 indicates that the drugs are synergistic, additive or antagonistic, respectively. In addition, we employed Bliss and Loewe models and SynergyFinder+, an open-source package (50), which can be accessible at www.synergyfinderplus.org, to quantify experimental data for drug synergy.

To comparatively assess the effect of the single-drug and combination treatments, we introduced a theoretical cell viability (ICV) function, defined as the aggregate of the activated levels of the major pro-growth signalling nodes as follows:

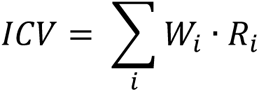

where *R_i_* is the level of activated pro-growth signalling proteins (pAKT, pERK, pS6K) and *W_i_* is the corresponding protein’s contributing weight to cell viability (see Supplementary Text 1 for more detail).

### Data availability

The data generated in this study are available within the article and its supplementary data files, and raw data can be provided upon request from the corresponding authors.

## Acknowledgements and Funding

L.K.N was supported by a Research Fellowship from Victorian Cancer Agency (MCRF18026). L.K.N’s laboratory was supported for this work by an Investigator Initiated Research Scheme grant from National Breast Cancer Foundation (IIRS-20–094) and a Venture Grant from Cancer Council Victoria. R.J.D’s laboratory was supported for this work by a Venture Grant from Cancer Council Victoria.

## Author Contributions

Conception and design: R.J.D, L.K.N.

Development of methodology: SY.S., N.J.C., M.G., L.K.N, R.J.D.

Acquisition of data: N.J.C., A.C.C., SY.S., M.G., YH.J.

Modelling and simulation: SY.S., M.G.

Analysis and interpretation of data: SY.S., N.J.C., M.G., A.C.C., YH.J., L.K.N, R.J.D.

Writing of the manuscript: SY.S., N.J.C., L.K.N, R.J.D.

Study supervision: R.J.D, L.K.N.

All authors read and approved the final manuscript.

