## Supplementary Information for "Integrative Modelling of Signalling Network Dynamics Identifies Cell Type-selective Therapeutic Strategies for FGFR4-driven Cancers"

### SUPPLEMENTARY MATERIALS

#### S1. Model assumptions, implementation and calibration

##### *Detailed description of the model scope and assumptions*

This model includes the major effector signalling pathways downstream of FGFR4, such as the FRS2, PI3K/AKT/mTOR and Ras/RAF/MEK/ERK signalling pathways (schematic diagrams are given in Figs 2A and S2). Since the latter two pathways are also converged upon by other receptor tyrosine kinases (RTKs) beside FGFR4, the model also incorporates the RTKs of the IGF1R/IR and ErbB receptor families.

*Activation of RTKs and the downstream oncogenic pathways.* The four RTKs of the ErbB receptor family undergo both homotypic and heterotypic interactions and in order to capture this in a simple manner we have incorporated a generic ErbB heterodimer into our model (1). Activated IGF1R predominantly activates the PI3K-PDK1-AKT pathway but can also enhance RAS-ERK signalling via growth factor receptor bound 2 (Grb2) and son of sevenless (Sos). Upon activation, FGFR4 phosphorylates FRS2, a docking protein and intracellular target of the FGFR receptors. FRS2 activation results in recruitment of Grb2 and Sos, which can activate the Ras/RAF/MEK/ERK signalling pathway (2,3). Activation and phosphorylation of specific ErbBs recruit Grb2 (4), leading to activation of Shp2 and Sos, which activates Ras-Raf-MEK-ERK signalling and the PI3K-PDK1-AKT pathway through Gab1/2 (5). In our model, the ErbB heterodimer activates PI3K signalling either via Grb2/Gab or Ras. Although PI3K can also be activated by ErbBs through p85 binding to ErbB3 (6), this is omitted for simplification.

*Major feedback regulation within the network*. Grb2 binds to tyrosine-phosphorylated FRS2, and Grb2 also forms a complex with the E3 ligase Cbl by means of the Grb2 Src homology 3 domains, resulting in the ubiquitination of FGFR and FRS2 in response to FGF stimulation (7-9). Thus, this forms a negative feedback loop between Cbl and FGFR4. SPRY2 expression is induced by ERK activation (10,11). After cell stimulation by growth factors SPRY2 can translocate to bind to the scaffold protein FRS2 and become phosphorylated on a conserved tyrosine (Tyr55) (12,13), which provides docking sites for the adaptor protein Grb2 (13). SPRY2 binding to Grb2 inhibits the recruitment of the Grb2-Sos complex either to FRS2 or to Shp2 (14). Thus, we assumed in the model that phosphorylated SPRY2 inhibits activation of Grb2/Ras. In addition, SPRY2 also inhibits the Ras/ERK pathway by supressing the activation of Raf (15) through a negative feedback loop, which was also incorporated in our model.

The signalling activity of RTKs is determined not only by the kinase activity of the RTKs themselves but also by the activities of protein tyrosine phosphatases (PTPs), which can dephosphorylate RTKs on tyrosine residues and attenuate their signalling capacity (16,17). In this model we considered PTPs-mediated negative regulation of RTKs by assuming they dephosphorylate and inhibit the RTKs (18,19). Along with the PTPs negative regulation of RTKs, experimental evidence of positive regulation of PTPs' transcription by AKT and ERK signalling together provide supporting evidence for PTP-mediated negative feedback loops to RTKs. For example, our analysis of drug perturbation data obtained from the Connectivity Map (CMap) project (20,21) demonstrate that the gene expression of PTPN12 was significantly reduced over time by AKT inhibitor or ERK inhibitor treatment in multiple cell lines (Fig. S29), suggesting activation of these pathways promotes expression of specific PTPs. Thus, we included in the model PTPs-to-RTKs negative feedback loops. For simplicity we assumed that PTPs represent a set of specific protein tyrosine phosphatases with similar functions towards the RTKs (e.g. PTPN12) (17), and so modelled it as a single entity.

*Crosstalk between the Ras/ERK and PI3K/AKT signalling pathways*. ERK phosphorylates Gab1 at six serine/threonine residues (T312, S381, S454, T476, S581, S597) (22), which inhibit Gab1/PI3K association and thus suppress the activity of PI3K (23). ERK can also phosphorylate Gab2 and negatively regulates p85 recruitment (24). AKT constitutively associates with Gab2, phosphorylates Gab2 on a consensus phosphorylation site, Ser159 and inhibits Gab2 tyrosine phosphorylation. Thus, in the model we assumed that ERK inhibits both Gab1 and Gab2 and AKT inhibits Gab2 only. In addition, we assumed that Ras directly activates PI3K through interacting via an amino-terminal Ras-binding domain (RBD) (25). Further, AKT phosphorylates and inhibits Raf, which leads to inhibition of the Raf-MEK-ERK cascade (26,27), providing another crosstalk point between the two pathways.

#### *Construction and implementation of the FGFR4 network model*

The new FGFR4 network model was formulated using ordinary differential equations (ODEs). The rate equations and full set of ODEs are given in Supplementary Tables S1-2. A reduced model schematic displaying the major network interactions is given in Figure 2A, and a detailed model reaction diagram containing all the model reactions is provided in Figure S2. The full model ODEs, rate equations and the sets of best-fitted parameter values used for simulations are provided in Supplementary Data S2-4. The model was implemented and numerically simulated in MATLAB (The MathWorks. Inc. 2023a) and the IQM Tools (https://iqmtools.intiquan.com/). The IQM Tools provide an interface to the Sundials ODE solvers (https://computing.llnl.gov/projects/sundials), which is used to convert a standard IQM model into user-callable MEX file for high performance computer simulation. To facilitate model exchange, an exchangeable Systems Biology Markup Language (SBML) file of the model is provided as Supplementary Data S1. In addition, the code for the modelling has been deposited to Github and can be accessed at: https://github.com/NguyenLabNetworkModeling/FGFR4-Signaling-Network-Model.git.

#### *Model calibration*

The adequacy of a mathematical model is generally justified by its ability to recapitulate known experimental data, which is ensured through a process known as model fitting or calibration where unmeasured model parameters are numerically estimated so that model simulations fit the data. Parameter estimation was done by minimizing the following ‘objective function’ that quantifies the discrepancy between simulated values and corresponding experimental measurements (28):

$$J(\boldsymbol{p}) = \sum_{j=1}^{M}\sum_{i=1}^{N}\left(\frac{y_{j,i}^{D} - y_j(t_i,\boldsymbol{p})}{\sigma_{j,i}}\right)^2$$

where $M$ is the number of the given experimental data sets used for fitting and $N$ is the number of time points within each experimental data set. $y_j(t_i,\boldsymbol{p})$ represents the numerical solution for the model state variable $y_j$ evaluated at time $t_i$ and parameter set $\boldsymbol{p}$; while $y_{j,i}^{D}$ is the mean value of the corresponding data point at $t_i$ with the associated variance of measured data $\sigma_{j,i}$.

A Genetic Algorithm (GA) was used to optimize the objective function (29-31). This was done by using the Global Optimization Toolbox and the function *ga* in MATLAB. Selection rules select the individual solutions with the best fitness values (called 'elite solution') from the current population. The elite count was set to 5% of the population size. Crossover rules combine two parents to generate offspring for the next generation. The crossover faction was set at 0.8. Mutation rules apply random changes to individual parents to generate the population of the next generation. For the mutation rule, we generated a random number from a Gaussian distribution with mean 0 and standard deviation $\sigma_k$, which was applied to the individuals of the current generation. The standard deviation function ($\sigma_k$) is given by the recursive formula as follows:

$$\sigma_k = \sigma_{k-1}\left(1 - \frac{k}{G}\right),$$

where $k$ is the $k^{th}$ generation, $G$ is the number of generations, and $\sigma_0 = 1$.

To derive at the best fitted parameter set, we carried out repeated GA runs with population size of 2000 and the generation number set to 200. In this computation, we also changed the mutation and crossover rates and even the population size to escape from being trapped in local minima. After multiple repetitions of the GA process where the best fitted set obtained from a previous repeat was used as the starting point of the next repeat, we arrived at the final best fitted set as the objective function was not further reduced, and the fitted parameter values no longer change. This whole process was repeated multiple times to obtain 50 independent equally best-fitted parameter sets (Supplementary Data S2-4 and Figs S30, S32) which was collectively used for all simulations.

***Model identifiability analysis***

To convey the identifiability of the model parameters during model calibration, we have undertaken identifiability analysis for all the models using a well-established method based on *profile likelihood* (28,32,33). This method is able to detect both structural and practical non-identifiable parameters through calculating confidence intervals defined by a threshold in the profile likelihoods, as detailed in this method paper (33). A parameter is considered to be *identifiable* if its confidence interval of its estimate is finite, *practically non-identifiable* if the

confidence interval is semi-finite (c,∞) or (-∞, c), and *structurally non-identifiable* if the confidence interval is infinite (-∞,∞) (33). Non-identifiable parameters, both *practically* and *structurally*, indicate that the values of the parameters cannot be uniquely determined from the observed data, resulting in multiple possible parameter sets that yield similar calibration results.
For the parameter estimation we used the same objective function sed for the model calibration above. The profile likelihood of a parameter θ is given by (28,32,33)

$$\chi^2_{PL}(\theta_i) \ = \min_{\theta_{j\neq i}}[\ \chi^2(\theta)]$$

Profile likelihood-based confidence interval (CI) can be derived via

$$CI(\theta) = \{\chi^2_{PL}(\theta) - \chi^2_{PL}(\hat{\theta}) < \Delta_\alpha\},$$

where $\Delta_\alpha = \chi^2(\alpha, df)$ is the threshold, $\alpha$ is a confidence level (the α quantile of the $\chi^2$-distribution), $df$ is the degree of freedom ($df = 1$ for point-wise confidence interval and $df = \#\ of\ parameters$ for simultaneous confidence intervals, respectively). $\hat{\theta}$ denotes the best-fitted parameter set.

Figures S3-4, S12-13, and S18-19 display the identifiability analysis results for Model-1, -2, and -3, respectively. In each plot the black dash lines depict the cut-off threshold for the confidence interval. The results show for each parameter in a model whether it is *structurally* (indicated by a flat curve in both directions (33)) or *practically* (indicated by a flat curve only in one direction (33)) non-identifiable. Overall, these results demonstrate that all the models are non-identifiable, meaning each model has at least one non-identifiable parameter. As most large ODE-based models in systems biology are unidentifiable to some extent (34,35), the results here were not surprising given the detailed scope of our models, which were purposely designed to capture the important biological mechanisms within the FGFR signalling network, including multiple feedback/feedforward loops and crosstalk. Since there is a general trade-off between model identifiability and level of biological details, although the models in this study could be made more identifiable through model abstraction, such process is inevitably at the cost of sacrificing specific biological details and may weaken the models' explanatory and predictive power.

***Overall workflow of the model calibration process***

The identifiability analysis confirmed that the model is not identifiable, which was not surprising given the detailed scope of the model. Making the model completely identifiable is practically infeasible without compromising the level of detail in the model. Thus, we aimed to improve the model constrain and identifiability by reimplementing the model calibration-validation process. Specifically, instead of calibrating (fitting) the model once using the data, the model fitting process is repeated through three sequential phases, where at each phase new data generated to validate new model predictions are then incorporated into the pool of training data and the entire pool was used to recalibrate the model at the next phase, which was then used to make new sets of predictions. In simple terms, the model gradually 'evolved' overtime as it was increasingly refined and constrained in light of new experimental data. This implementation better reflects the iterative modelling-experimental cycles, which is the hallmark of the systems approach by our study.

The specific modelling workflow is given in Fig S31. Model-1 of the phase-1 uses the data in Fig 1A,D for model calibration. These data plus the time-course and dose-response data in response to PI3K inhibition (Fig S10) were used to recalibrate the model, generating Model-2 in the phase-2 (Fig S11). Finally, these data plus the time-course response data of specific RTKs (Fig 4B) and Fig S16 were used for model recalibration to generate a Model-3 in the phase-3 (Fig S17). In particular, to further constrain and improve the prediction accuracy, we additionally used time-course response data to ERBB and AKT inhibitors (Fig S17). Note the time-course of ERBB inhibitor was reproduced from previously published data (36)

***Landscape of the estimated parameter values***

Through the model calibration process, we obtained 50 best-fitted parameter sets, which were collectively used for ensemble modelling and simulations. Each best-fitted parameter set was derived from independent re-start of the genetic algorithm (GA) with randomly sampled starting parameter values (within defined ranges) to ensure that the parameter space was comprehensively explored by the optimisation procedure.

The values for kinetic parameters associated with certain biochemical reactions can vary widely depending on the context. For instance, *Km* values, which represent the substrate concentration at which the enzyme's reaction rate is half-maximal, can range from nanomolar to millimolar concentrations (37-39). *Vmax* values, the maximum velocity of an enzyme-catalyzed reaction, can range from nanomoles to millimoles per minute (37-39). *kcat* values, the turnover number of an enzyme, can also vary widely, with some enzymes having *kcat* values in the range of a few per second to several hundred per second though the value of *kcat* is significantly influenced by factors such as the active site structure of the enzyme, the affinity of the enzyme for its substrate, and the stability of the enzyme-substrate complex (37,38,40). In order to account for this wide range of values, we have assumed the parameter range between $10^{-5}$ to $10^{5}$ for the kinetic parameter estimation (39,41,42). The estimated kinetic parameters, *Km*, *Vmax* and *kcat* are all in physically reasonable range on average (Fig. S32). However, it is important to be aware that the kinetic parameters of real biochemical reactions and mathematical models do not perfectly correlate in a technical sense because the model abstracts chemical reactions to a large extent.

***Simulation and computation of drug synergy***

The simulation and computation of drug synergy consists of four steps. The first step involves the simulation of dose-response against individual drugs. We gradually increased the concentration of the drugs and simulate the levels of the response readouts, often at a time point where steady state is reached after drug treatment. The second step is to estimate the IC50 values since the half-maximal inhibitory concentration (IC50) is a commonly used value for quantifying the potency of a drug and is often employed in drug combination studies. The IC50 value for each drug inhibitor was computed based on the dose-response curves from the previous step by fitting the dose-response curves to the four-parameter dose-response function (43) as follows:

$$Response = Bottom + \frac{Top - Bottom}{1 + \left(\frac{IC50}{X}\right)^{HillSlop}}$$

Where *Top* is the maximal value of response curve, *Bottom* is the minimal value (i.e., maximally inhibited response). *Top* and *Bottom* are plateaus of the curve in the units of the Y axis. *HillSlope* denotes the steepness of the sigmoidal curve. When *HillSlope* = 1, the dose-response is 'standard'; when HillSlope <1, the curve become 'shallower'; when HillSlope >1, the curve become 'steeper'. Having determined the suitable dose for each drug (e.g. IC50), in the step three, the effect of the pair-wise drug combinations is be systematically simulated and compared, called a 'dose-response matrix' setup. A typical setup involves combination of gradually increasing doses of two drugs A and B, around their respective IC50 values, including single-drug treatments. For example, the drugs could be combined at five doses: 1/5, 1/2, 1, 2 and 5 folds of the IC50 concentration. The last step is to quantify possible synergy (or lack thereof) between the individual drugs. To this end, different approaches can be used to compute drug synergism, depending on the specific context and preference. In this simulation study, we used Coefficient of Drug Interaction (CDI) (44,45) and highest single agent (HSA) (46) as a reference model. In addition, we employed Bliss and Loewe models and SynergyFinder+, an open-source package (47), which can be accessible at www.synergyfinderplus.org, to quantify experimental data for drug synergy.

**Model-based simulation and computation of drug synergy**

Because the synergy between two drugs can depend on the specific doses at which they are combined, we used IC50 value of individual drugs as a reference concentration for the simulation (see Supplementary Information for details). Using the simulated dose-response matrix, we evaluated in silico the efficacy and possible synergism of 19 possible combinatorial strategies co-targeting FGFR4 and each of 19 network components. Drug synergy was computed based on the coefficient of drug interaction (CDI) and highest single agent (HSA) metric (44-46): CDI = $R_{12}/(R_1 \times R_2)$, where $R_{12}$ is a normalized biological response (e.g., cell viability) by the combined treatment of drug 1 and 2, and $R_1$ and $R_2$ are the response by the single drug treatment, respectively. Note $R$ = 1 (and 0) indicates no drug response (complete inhibition, respectively). CDI <1, = 1 or >1 indicates that the drugs are synergistic, additive or antagonistic, respectively. According to the HSA model, the synergistic effect is equal to the greater effect of individual drugs (46). Thus, HSA = $E_{12} - \max(E_1, E_2)$. $E_{12}$ is a normalized drug effect by the combined treatment of drug 1 and 2, and $E_1$ and $E_2$ are the response by the single drug treatment, respectively. Note $E$ = 0 (and 1) indicates no drug effect (maximum effect, respectively). For the readability and easy comparison between CDI and HAS, we used a log2-transformed CDI metric (-log2(CDI)): Thus, for the both metrics the score > 0, = 0 or < 0 indicates that the drugs are synergistic, additive or antagonistic, respectively. In addition, we employed Bliss and Loewe models and SynergyFinder+, an open-source package (47), which can be accessible at www.synergyfinderplus.org, to quantify experimental data for drug synergy.

We comparatively assessed the effect of the single-drug and combination treatments by introducing a theoretical cell viability (ICV) function, defined as the aggregate of the activated levels of the major pro-growth signalling nodes as follows:

$$ICV = \sum_i W_i \cdot R_i$$

where $R_i$ is the level of activated pro-growth signalling proteins and $W_i$ is the corresponding protein's contributing weight to cell viability.

In this study, we considered the three readouts pAKT, pERK, pS6K as pro-growth marker proteins since these represent the outputs of the three key downstream signalling pathways in our model, the PI3K/AKT, Ras/ERK and mTOR/S6K pathways that are converged upon by the upstream RTKs, and that these readouts are well-established markers of cell proliferation and survival. Given that the contribution of the three oncoproteins to cell viability are different, we utilized drug response data directed at these nodes measured in the MDA-MB-453 cells to inform the relative weights of each readout. For example, because complete inhibition of AKT using high dose of a specific AKT inhibitor (MK-2206) potently blocked MDA-MB-453 viability by ~97%, while complete inhibition of ERK only blocked MDA-MB-453 viability by 41%, this suggests pAKT plays a relatively more dominant role than pERK in driving cell viability. Accordingly, we used the maximal drug effect to inform the weight for the readouts, as shown in Supplementary Table S4, and employed the weighted sum to describe the final ICV function.

**S2. Supplementary Figures**

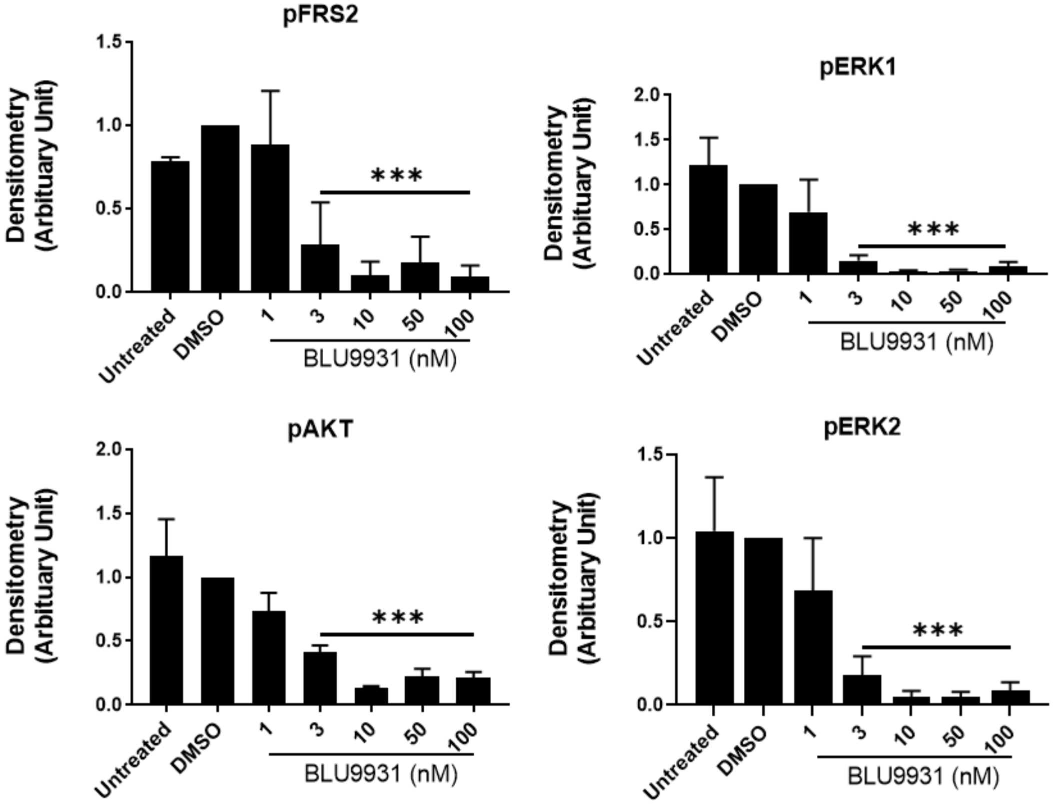


**Figure S1.** Quantification of dose dependent effect of BLU9931 in MDA-MB-453 cell line on expression and activation of downstream signalling proteins 1 h post-treatment with the indicated doses (see Figure 1A). *** indicates p-value of < 0.001, comparing individual BLU9931 concentrations to the DMSO vehicle control.

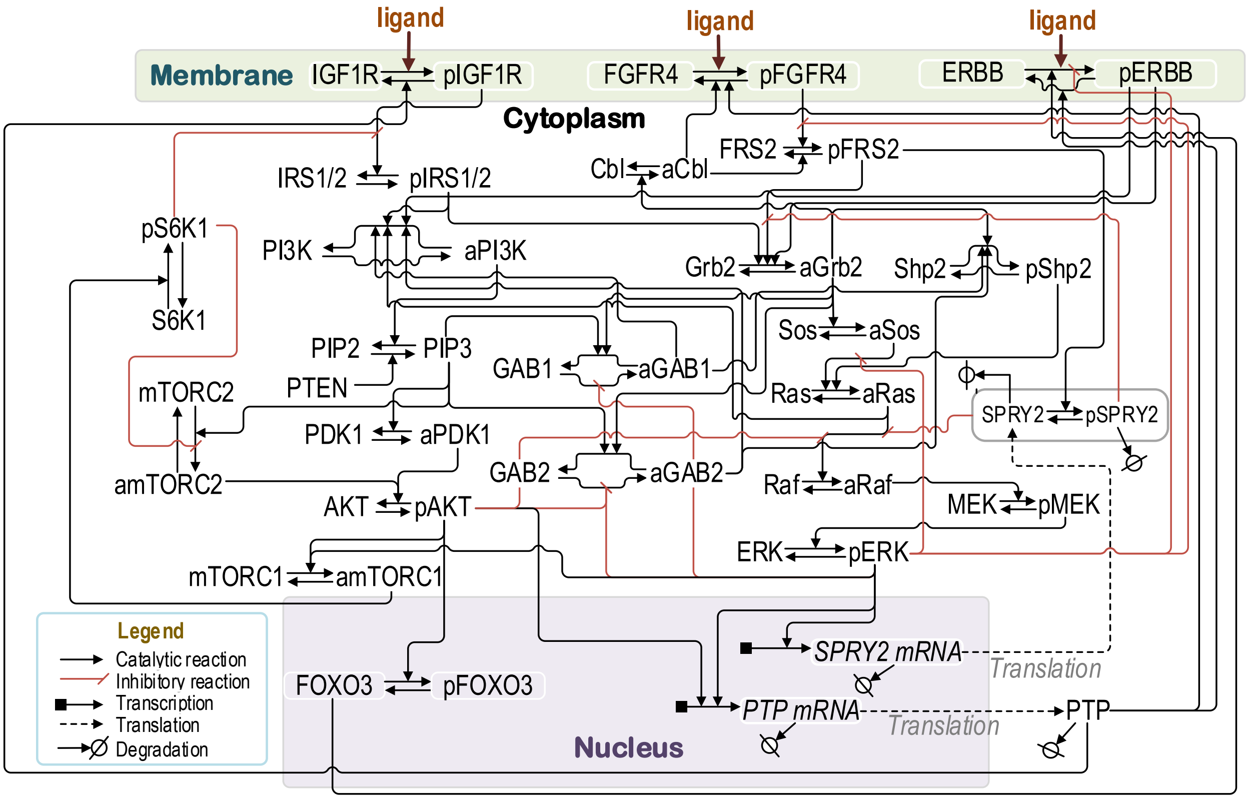


**Figure S2. Detailed reaction scheme of the FGFR4 network model.** This schematic diagram displays the detailed biochemical reactions included in the model. The reaction rates and model ODEs are given in Table S1-2.

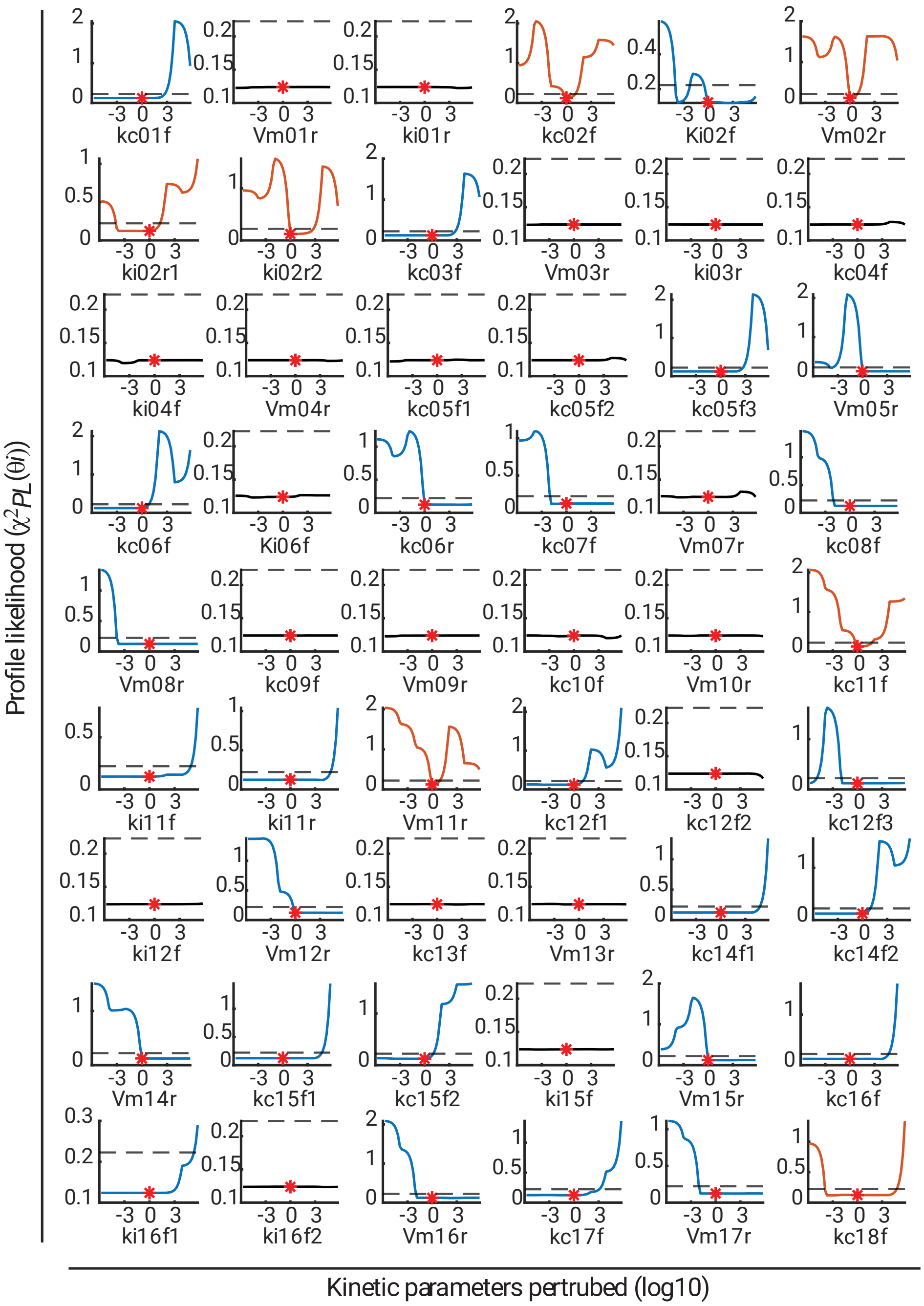


**Figure S3. Parameter identifiability analysis (Model-1, Part A)**. The profile likelihood for each parameter is displayed versus the model parameters. The red star at the lowest point for each parameter indicates the best-fitted parameter value. The relative value of the parameter compared to the best-calibrated value is depicted on the x-axis. The black dashed line denotes the cut-off threshold for the confidence interval. The identifiability of each parameter is represented by the colour of the curve: red, blue and black signifies *identifiable*, *practically non-identifiable,* and *structurally non-identifiable*, respectively.

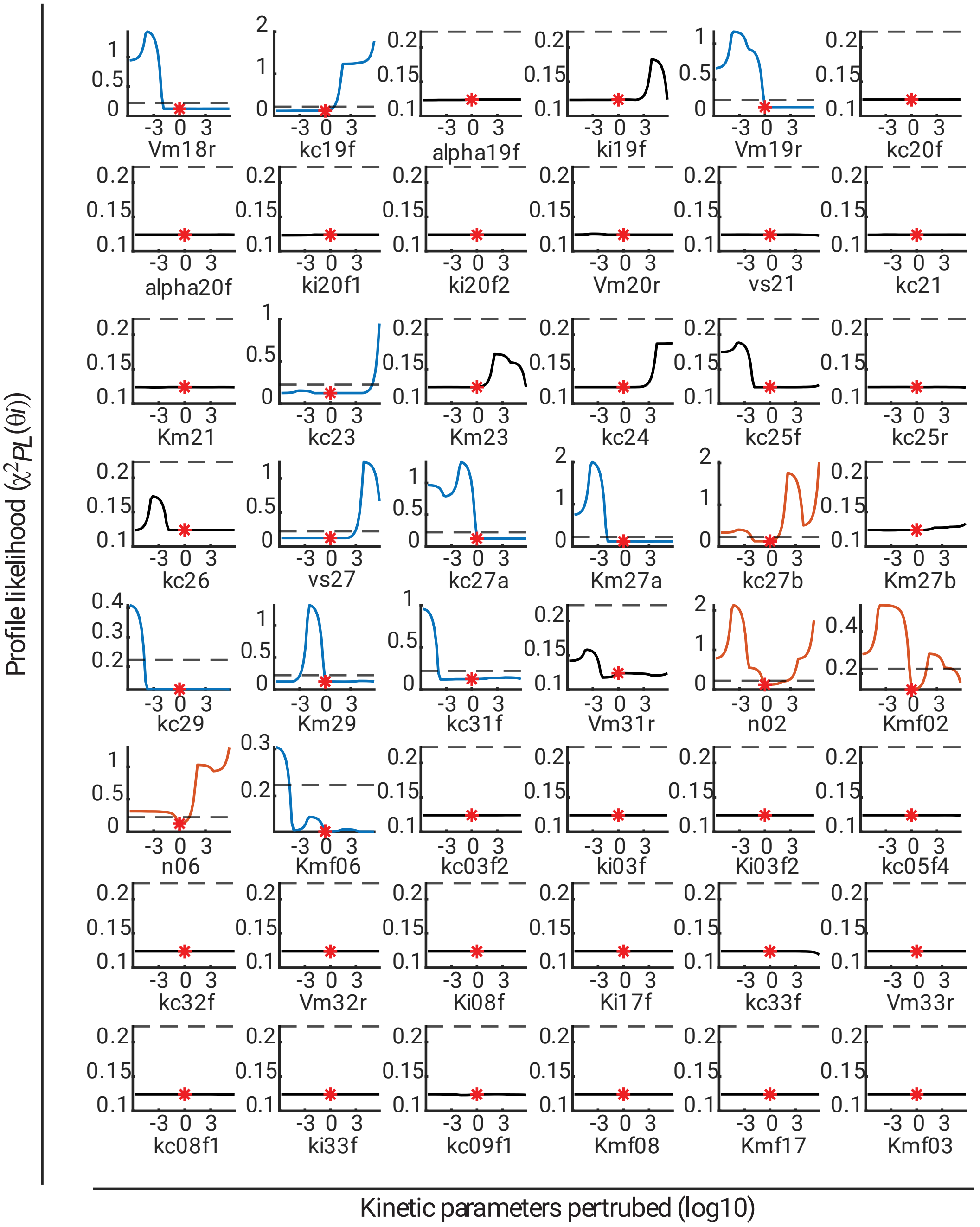


**Figure S4. Parameter identifiability analysis (Model-1, Part B)**. The profile likelihood for each parameter is displayed versus the model parameters. The red star at the lowest point for each parameter indicates the best-fitted parameter value. The relative value of the parameter compared to the best-calibrated value is depicted on the x-axis. The black dashed line denotes the cut-off threshold for the confidence interval. The identifiability of each parameter is represented by the colour of the curve: red, blue and black signifies *identifiable*, *practically non-identifiable*, and *structurally non-identifiable*, respectively.

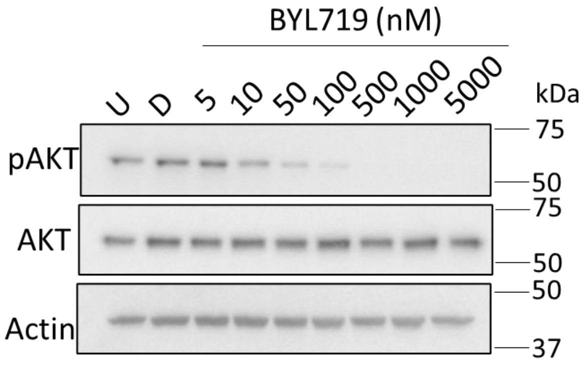


**Figure S5. Dose dependent effect of PI3Kα inhibitor BYL719 on AKT phosphorylation in the MDA-MB-453 cell line.** Expression and activation of AKT after 1 h treatment with BYL719 (representative of three independent replicates).

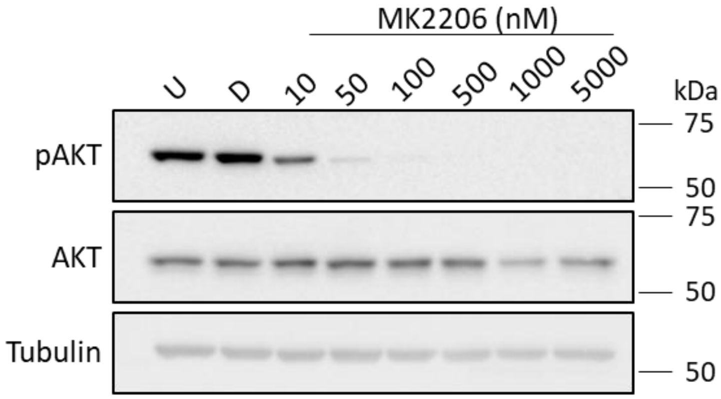


**Figure S6. Dose dependent effect of AKT inhibitor MK2206 on AKT phosphorylation in the MDA-MB-453 cell line.** Expression and activation of AKT after 1 h treatment with MK2206. U indicates untreated control, D indicates DMSO vehicle control. Representative of three biological replicates.

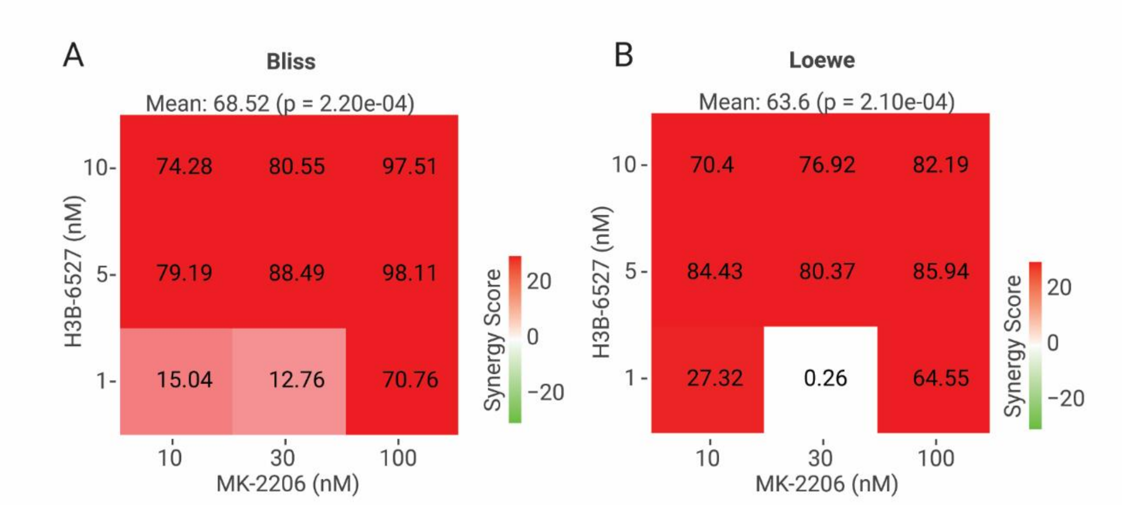


**Figure S7. Quantitative assessment of drug synergism.** Quantitative assessment of drug synergism between H3B-6537 and MK-2206 in the MDA-MB-453 cell line using Bliss score (A) and Loewe score (B). To calculate the synergy score, we used SynergyFinder+, an open-source package (47), which can be accessible at www.synergyfinderplus.org.

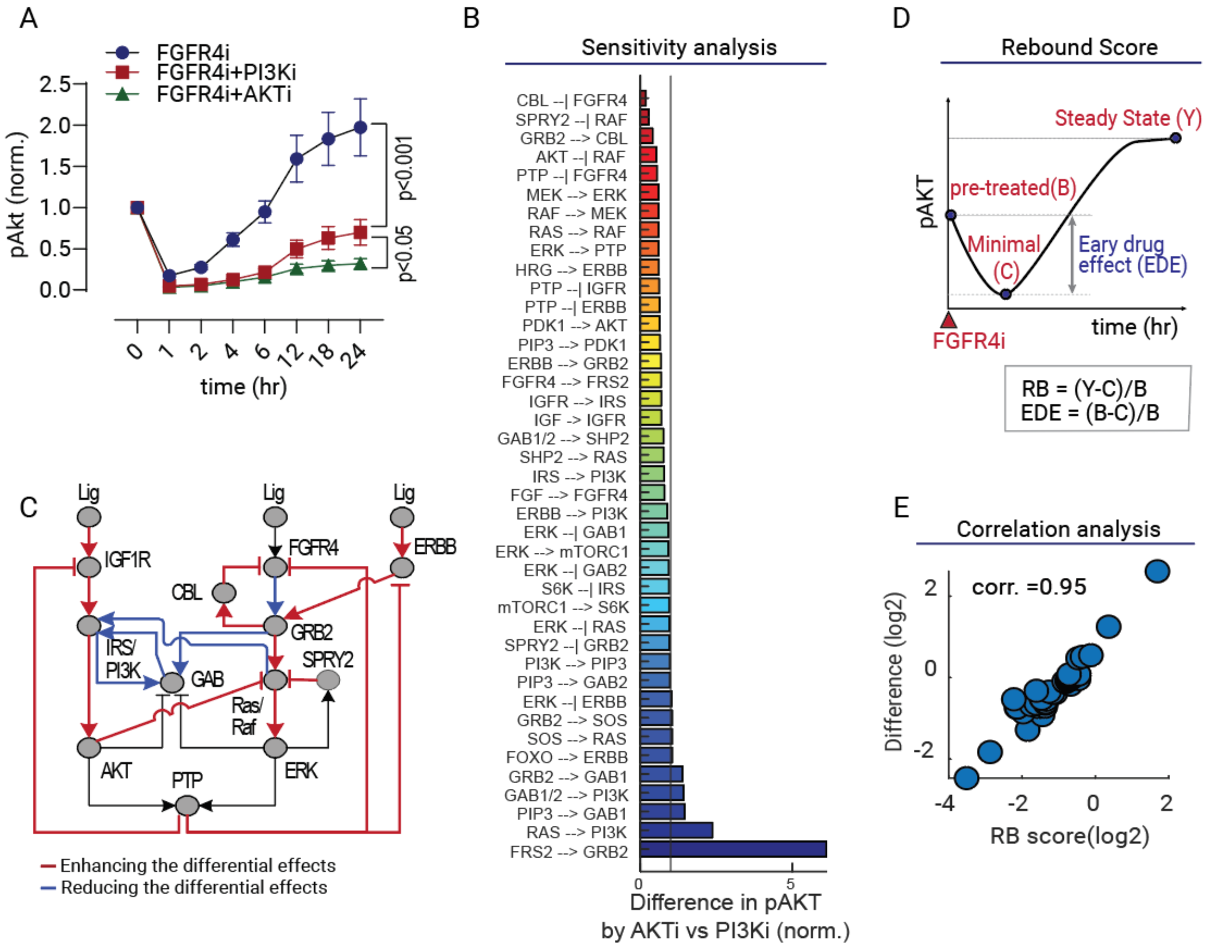


**Figure S8. Differential impact of combined FGFR4 with AKT vs. PI3K targeting on pAKT rebound dynamics analysed via model simulations. (A)** Simulated dynamics of phosphorylated AKT (pAKT) after FGFR4 inhibition (FGFR4i) alone or in combination with PI3K inhibition (PI3Ki) or AKT inhibition (AKTi). Each target was inhibited at its respective IC90 concentration. **(B)** Sensitivity analysis showing the differential effects of AKTi and PI3Ki on FGFR4i-induced pAKT rebound following specific network perturbations. Kinetic parameters representing network connections were individually blocked (by 99%), and the impact on pAKT levels at 24 hours under AKTi and PI3Ki was quantified, normalized to control conditions. A score (y-axis) <1, =1, or >1 indicates decreased, unchanged, or increased differential suppression of pAKT rebound by AKTi compared to PI3Ki. Averages were calculated using all 50 best-fitted parameter sets. **(C)** Simplified network diagram highlighting key links with significant impact from the sensitivity analysis (B). **(D)** Illustration of pAKT rebound dynamics post-FGFR4 inhibition and associated quantitative metrics, including Rebound (RB) and Early Drug Effect (EDE) scores. Higher RB scores signify stronger pAKT rebound; higher EDE scores indicate more pronounced initial drug (FGFR4i) impact on pAKT. **(E)** Scatter plot illustrating a strong positive Pearson correlation between pAKT RB score and the steady-state pAKT difference under AKTi vs. PI3Ki treatment. Each point represents the RB score and differential effect between PI3Ki and AKTi for perturbed network links. This analysis is based on Figure S9, indicating that greater pAKT rebound magnitudes enhance the differential effect of AKTi over PI3Ki.

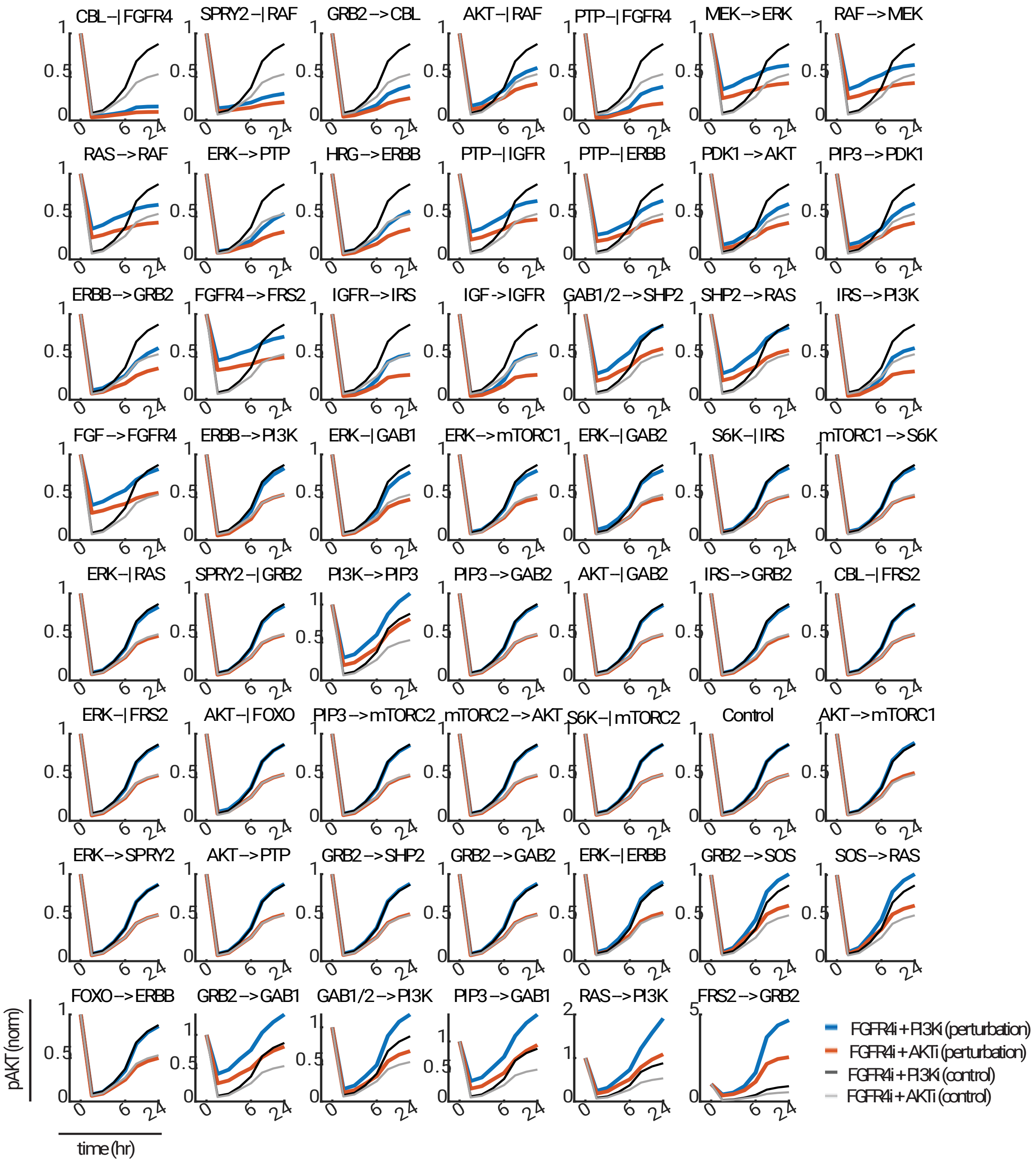


**Figure S9. Time-course simulations of pAKT dynamics for the sensitivity analysis in Fig S8B.** The dynamic responses of pAKT following co-inhibition of FGFR4 and either AKT or PI3K were simulated, comparing control condition (grey lines) and when each of the indicated links was blocked by 99% (blue/red lines). All the 50 best-fitted parameter sets were used for model simulations, and the results were averaged.

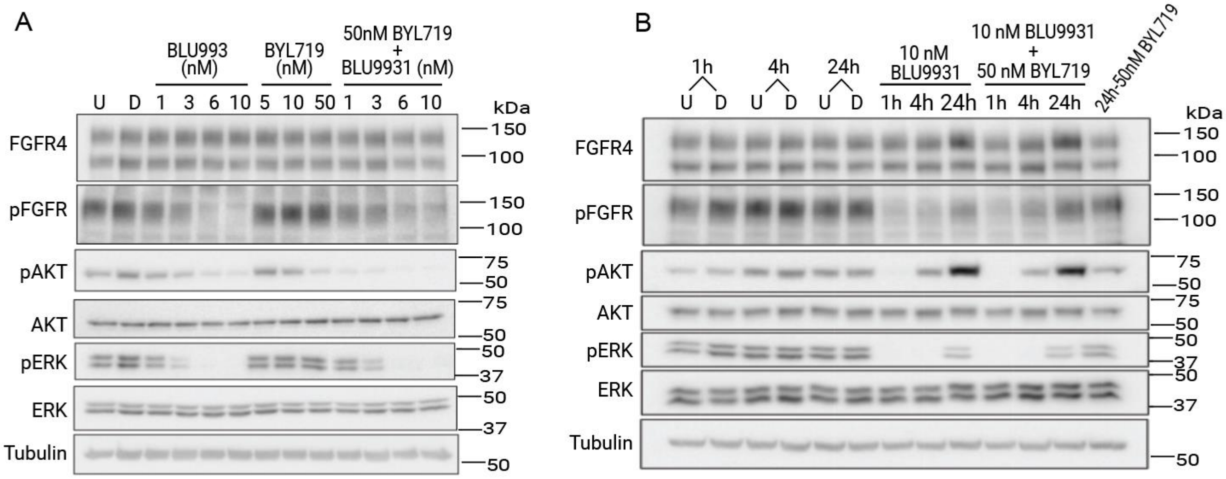


**Figure S10. Combined effect of the FGFR4 inhibitor BLU9931 and PI3K inhibitor BYL719 on FGFR4 downstream signalling pathways in the MDA-MB-453 cell line. (A)** Dose-dependent effect of BLU9931 and BYL719 on activation of downstream signalling proteins in the MDA-MB-453 cell line 1 h post-treatment with the indicated doses individually and in combination. (**B**) Time course analysis of combined treatment of BLU9931 and BYL719 on FGFR4 downstream signalling pathways in the MDA-MB-453 cell line. Expression and activation of downstream signalling proteins 1, 4 and 24 h post-treatment with 10 nM of BLU9931 and 10 nM of BLU9931 + 50 nM of BYL719, respectively. Note that the pFGFR signal was detected using a pan-phospho-FGFR antibody.

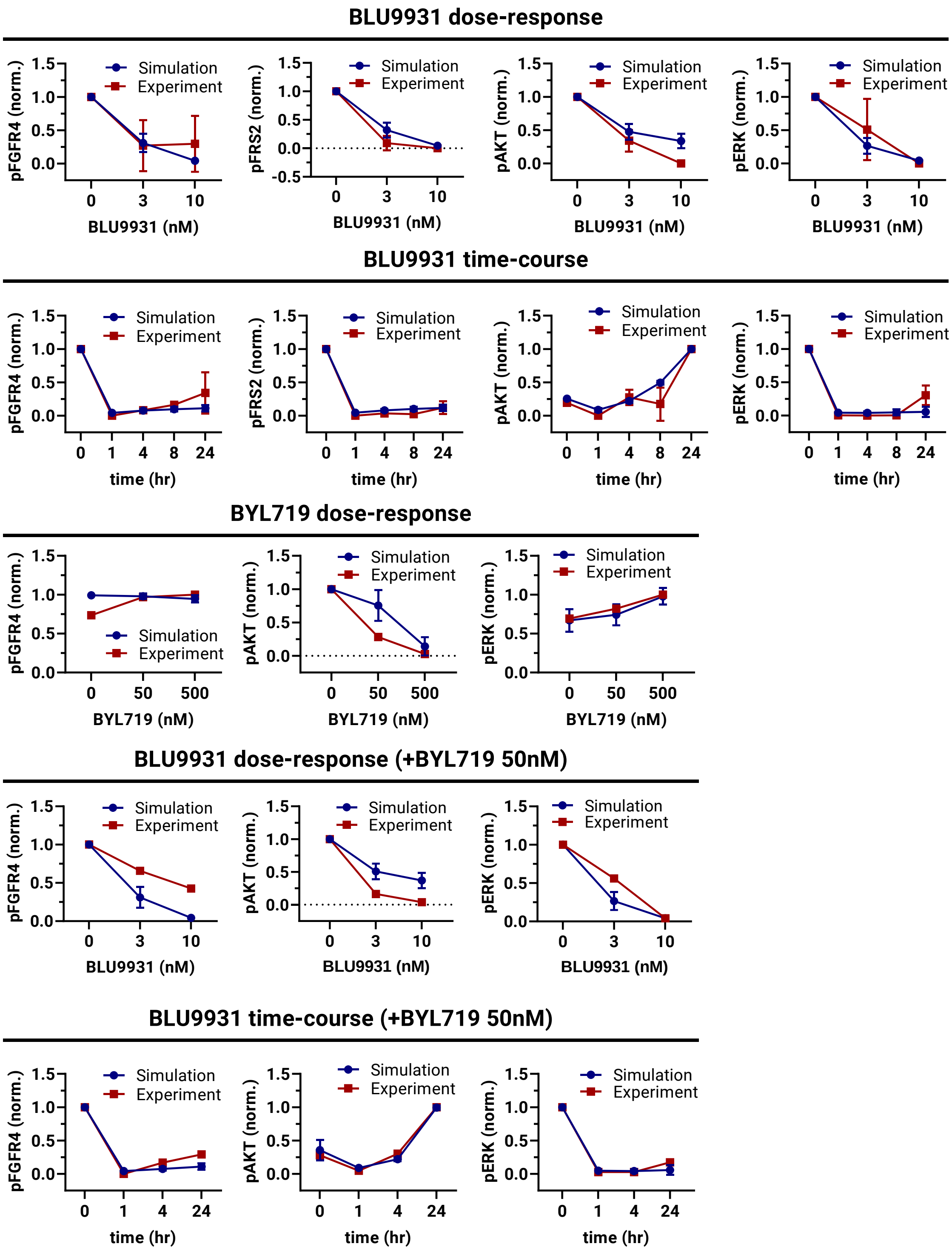


**Figure S11. Model calibration results of Model-2 (phase-2).** Comparison of model simulation using best-fitted parameter sets (blue lines, error bars: mean ± standard error, n=50) against experimental time-course and dose-response data (red lines, error bars: mean ± standard error, n≥2) demonstrate good agreement between simulation and data. Data used for additional model calibration were quantified from Figure S10.

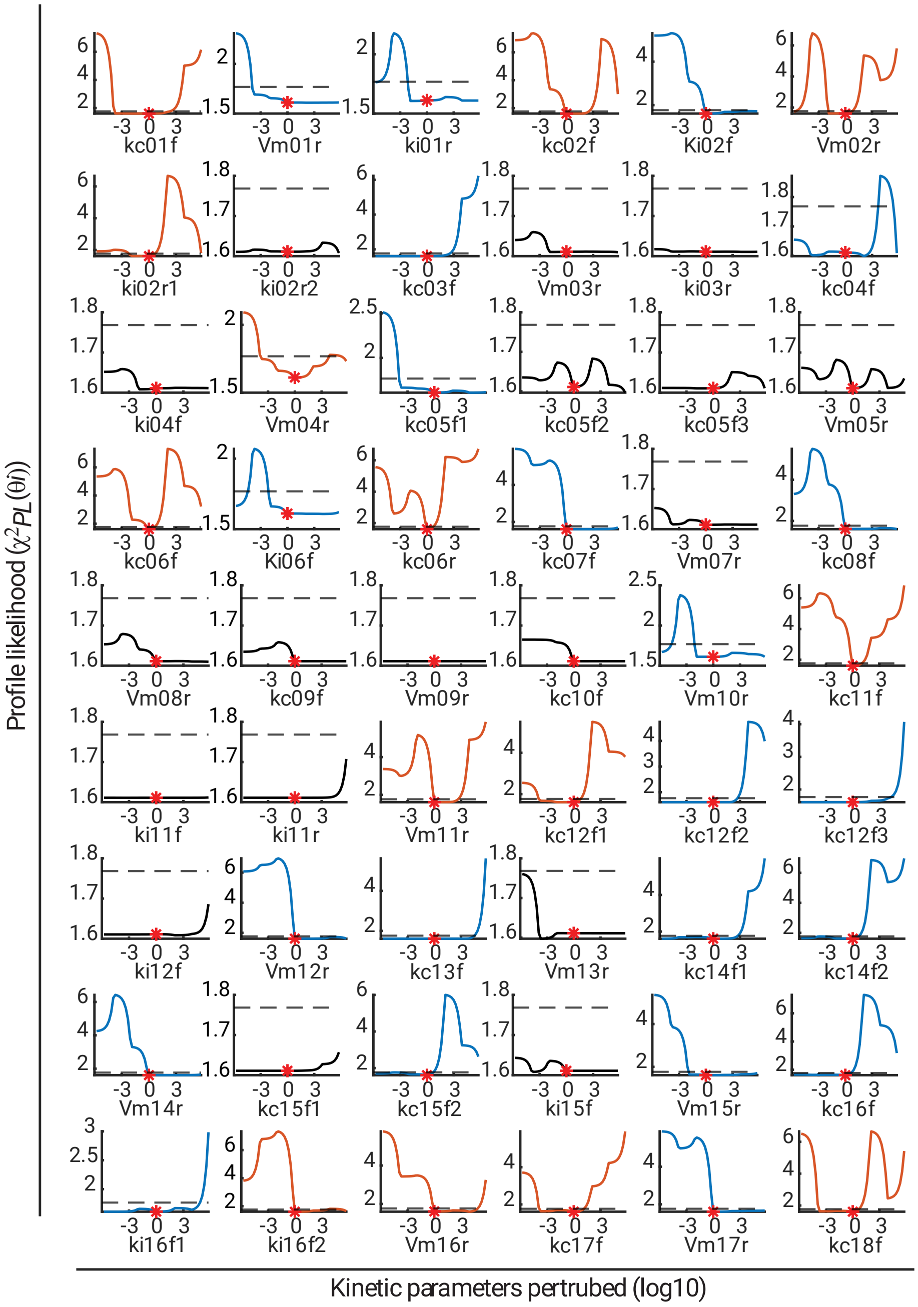


**Figure S12. Parameter identifiability analysis (Model-2, Part A)**. The profile likelihood for each parameter is displayed versus the model parameters. The red star at the lowest point for each parameter indicates the best-fitted parameter value. The relative value of the parameter compared to the best-calibrated value is depicted on the x-axis. The black dashed line denotes the cut-off threshold for the confidence interval. The identifiability of each parameter is represented by the colour of the curve: red, blue and black signifies *identifiable*, *practically non-identifiable,* and *structurally non-identifiable*, respectively.

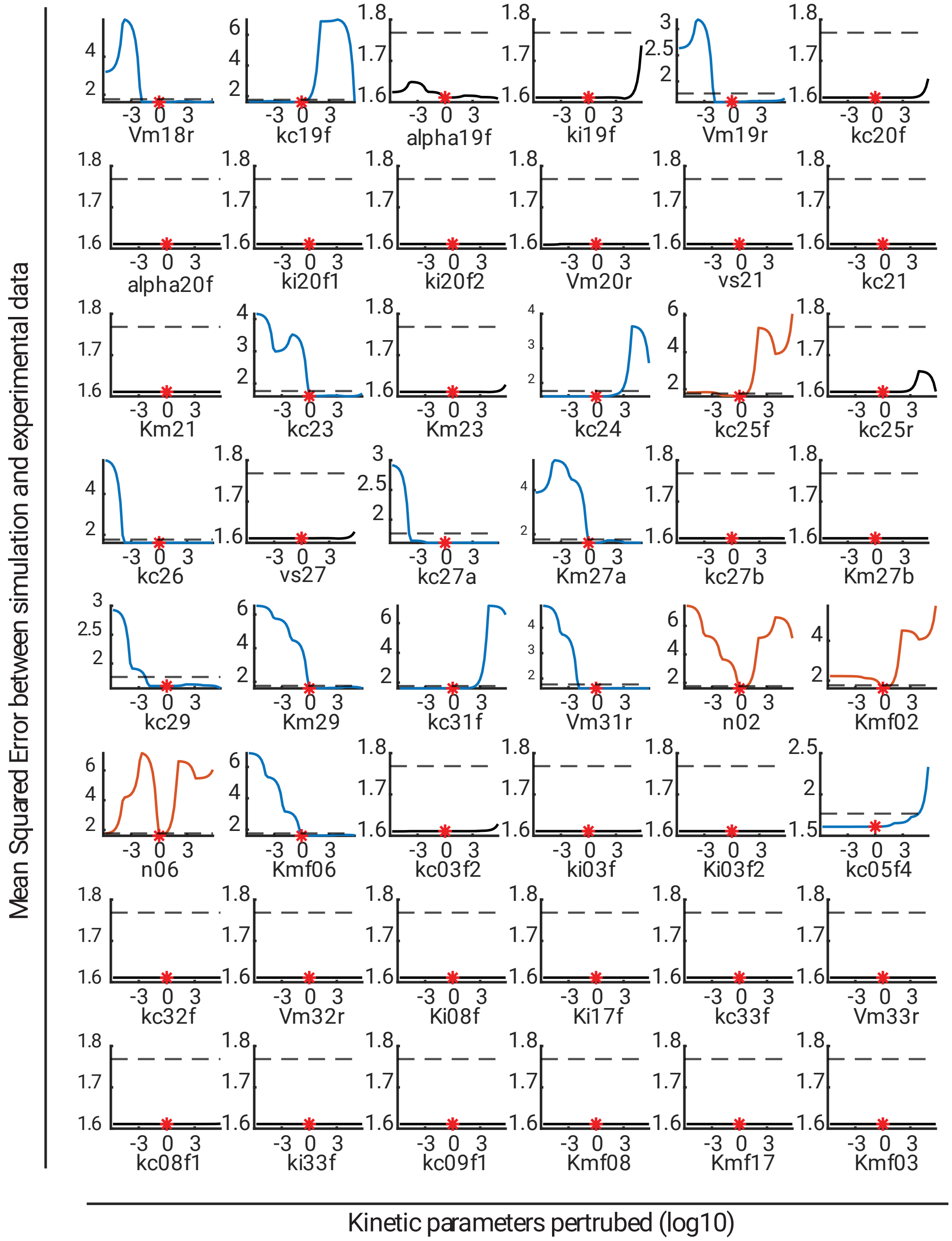


**Figure S13. Parameter identifiability analysis (Model-2, Part B)**. The profile likelihood for each parameter is displayed versus the model parameters. The red star at the lowest point for each parameter indicates the best-fitted parameter value. The relative value of the parameter compared to the best-calibrated value is depicted on the x-axis. The black dashed line denotes the cut-off threshold for the confidence interval. The identifiability of each parameter is represented by the colour of the curve: red, blue and black signifies *identifiable*, *practically non-identifiable*, and *structurally non-identifiable*, respectively.

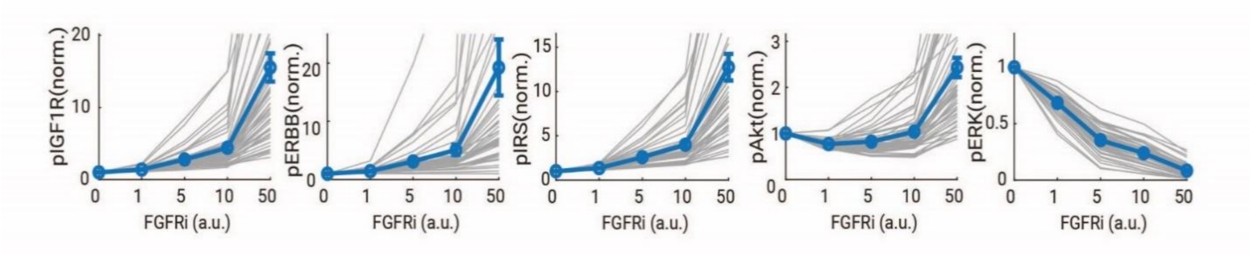


**Figure S14. Model simulation of the steady-state response of various network components to increasing BLU9931 treatment in MDA-MB-453 cells.** The thick blue line represents average of simulations using 50 best-fitted parameter sets. Error bars: mean ± standard error (n=50). The thin grey lines indicate simulation resulting from each individual best-fitted parameter set.

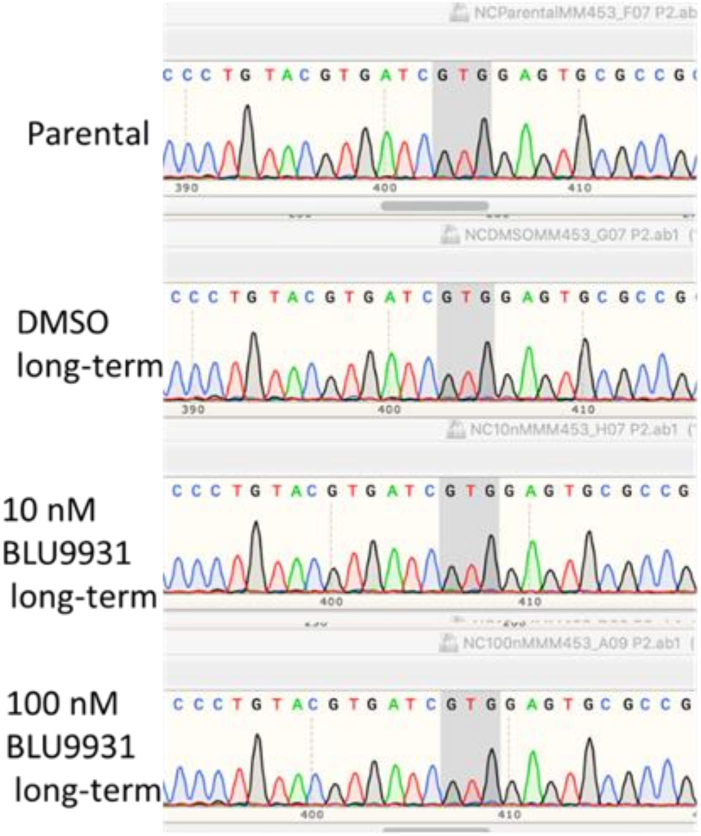


**Figure S15. FGFR4 kinase domain analysis for gatekeeper mutations.** RNA was extracted from parental and long-term FGFR4 inhibitor treated MDA-MB-453 cells followed by RT-PCR and Sanger sequencing analysis. If a FGFR4 gatekeeper mutation is present, the original GTG coding for valine in the DNA sequence ATCGTGGAGTGCGCC would have a base pair change, changing the amino acid. Example FGFR4 gatekeeper mutations are V550M/V550L.

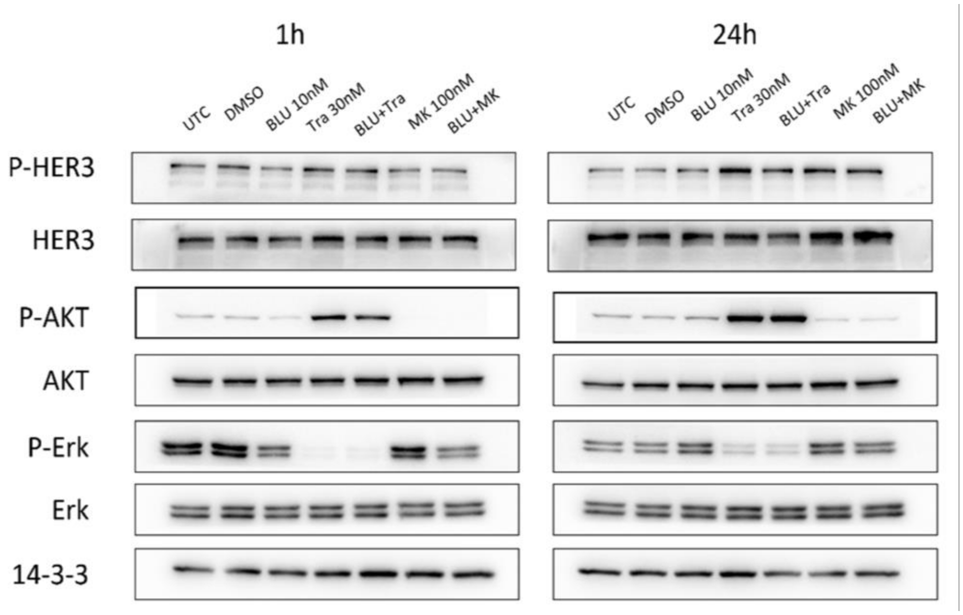


**Figure S16. Effect of AKT and MEK inhibitor treatment on FGFR4 downstream signalling pathways in the MDA-MB-453 cell line**. Expression and activation of downstream signalling proteins 1 and 24 h post-treatment with the indicated drugs. MK: MK-2206, BLU: BLU9931, Tra: Trametinib.

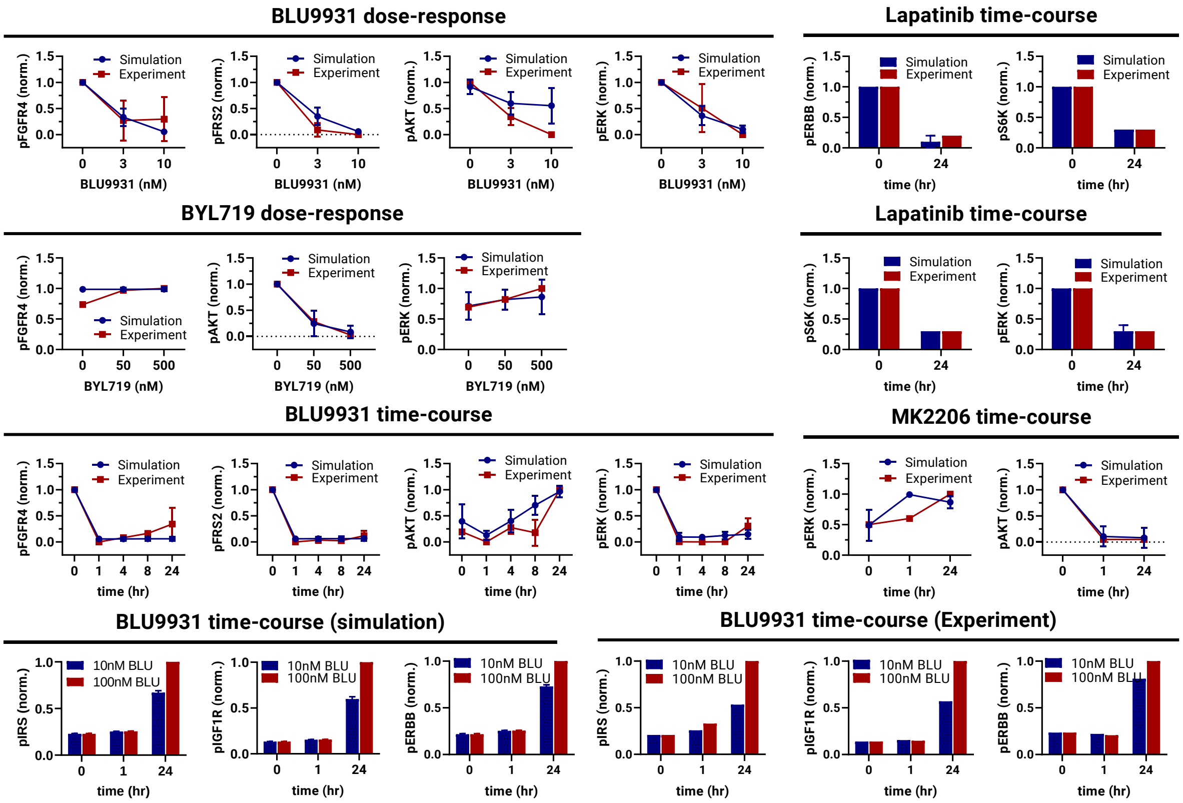


**Figure S17. Model calibration results of Model-3 (phase-3).** Comparison of model simulation using best-fitted parameter sets (blue lines, error bars: mean ± standard error, n=50) against experimental time-course and dose-response data (red lines, error bars: mean ± standard error, n≥2) demonstrate good agreement between simulation and data. Data used for additional model calibration were quantified from Fig S16. Lapatinib time-course data were reproduced from the previous paper (36).

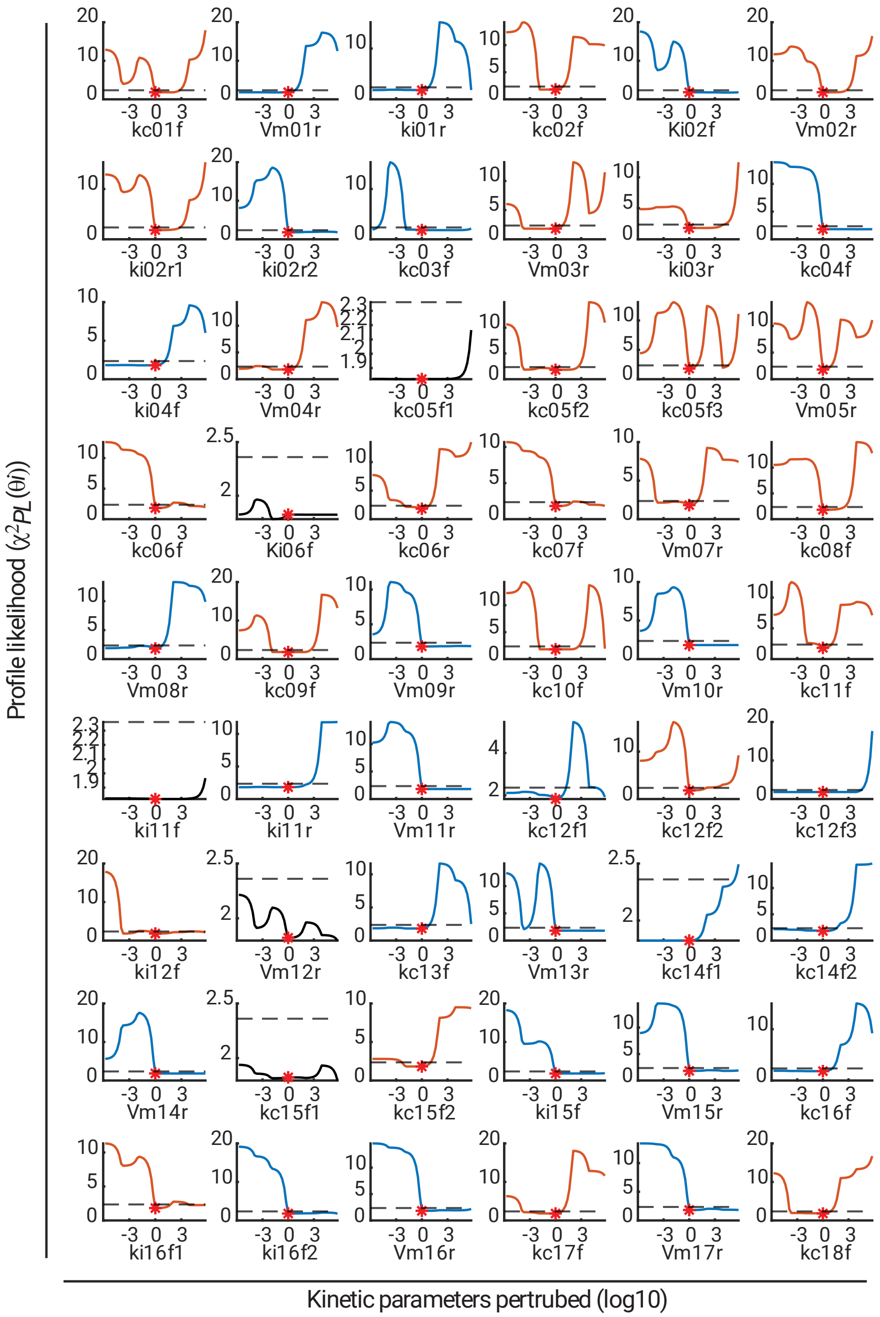


**Figure S18. Parameter identifiability analysis (Model-3, Part A)**. The profile likelihood for each parameter is displayed versus the model parameters. The red star at the lowest point for each parameter indicates the best-fitted parameter value. The relative value of the parameter compared to the best-calibrated value is depicted on the x-axis. The black dashed line denotes the cut-off threshold for the confidence interval. The identifiability of each parameter is represented by the colour of the curve: red, blue and black signifies *identifiable*, *practically non-identifiable,* and *structurally non-identifiable*, respectively.

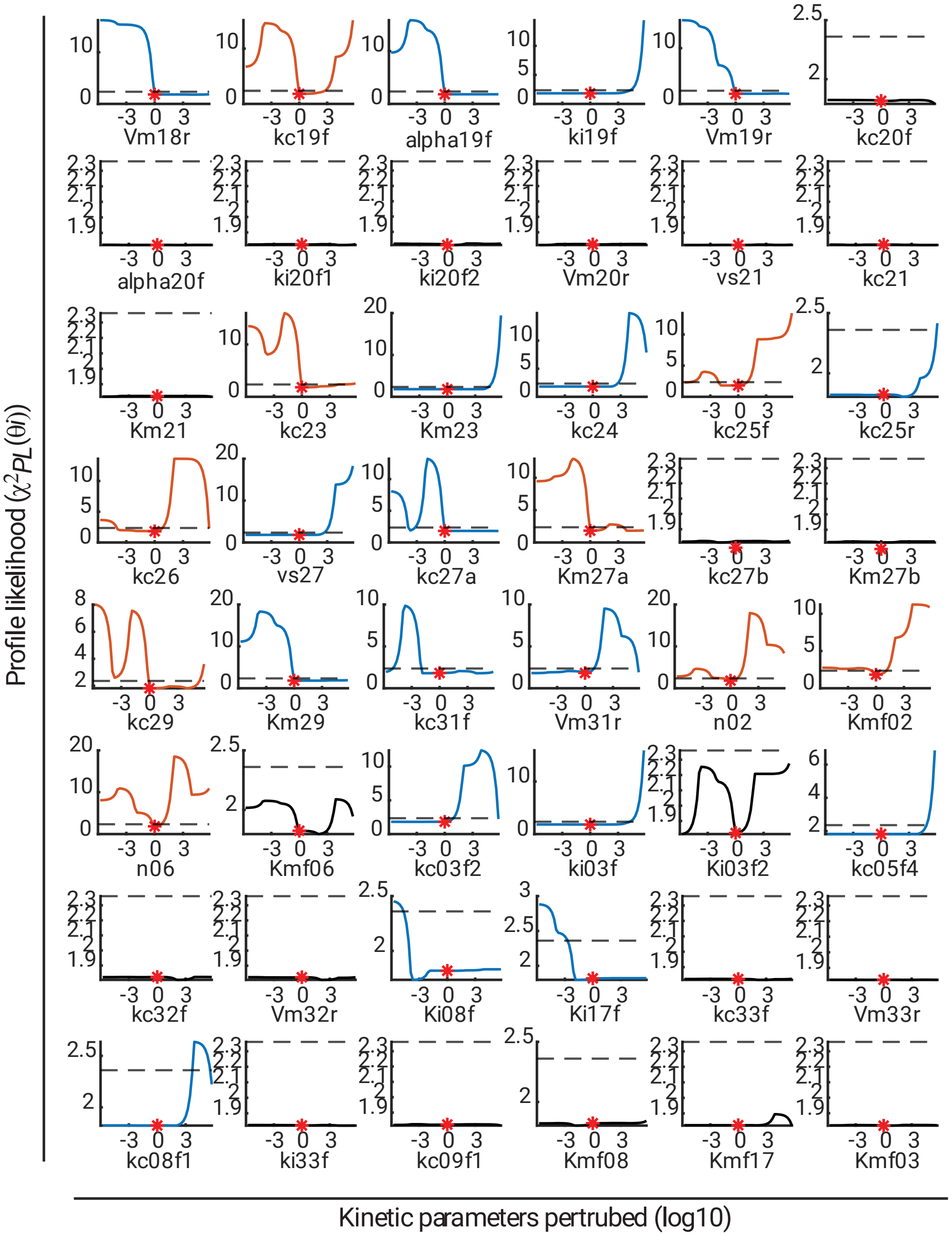


**Figure S19. Parameter identifiability analysis (Model-3, Part B)**. The profile likelihood for each parameter is displayed versus the model parameters. The red star at the lowest point for each parameter indicates the best-fitted parameter value. The relative value of the parameter compared to the best-calibrated value is depicted on the x-axis. The black dashed line denotes the cut-off threshold for the confidence interval. The identifiability of each parameter is represented by the colour of the curve: red, blue and black signifies *identifiable*, *practically non-identifiable*, and *structurally non-identifiable*, respectively.

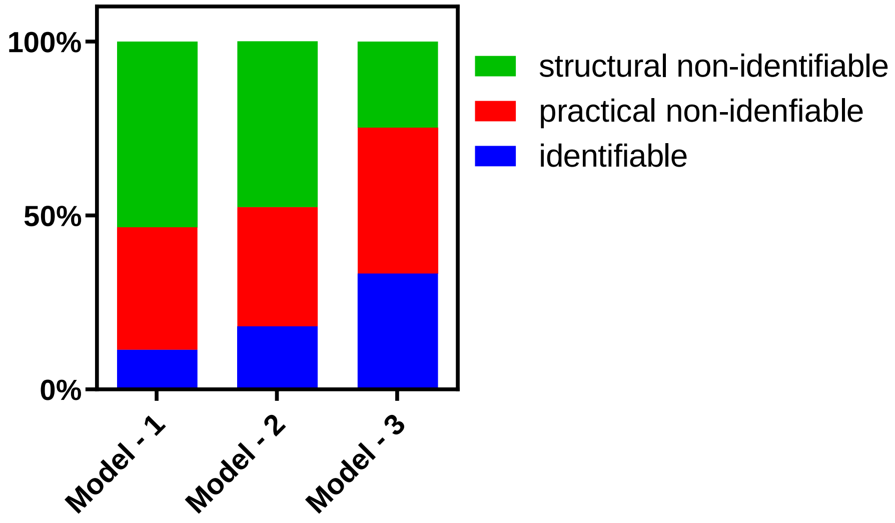


**Figure S20. Comparison of the parameter identifiability results for Model-1, -2 and -3**. The incorporation of additional training data sets into the model calibration process significantly increased the number of identifiable parameters, indicating an improvement in model constraint and ultimately leading to more precise and reliable model prediction results.

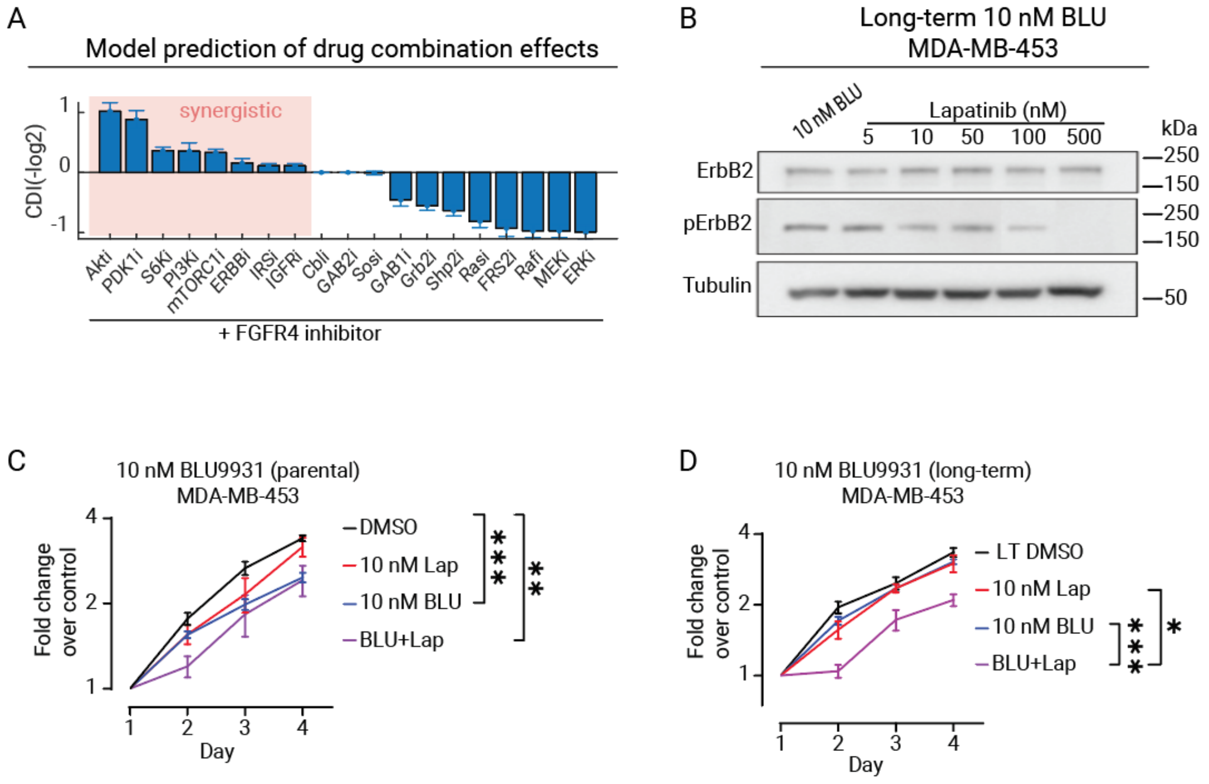

**Figure S21. Model prediction of synergistic drug combinations and experimental validation. (A)** In silico prediction of the effect of 19 possible pair-wise drug combinations co-targeting FGFR4 and various network components, assessed using drug synergy scores of the coefficient of drug interaction (CDI). The value that is >1, =1, <1 indicate synergistic, additive or antagonistic effects, respectively; and higher CDI values indicate stronger synergism. Error bars: mean ± standard error of 50 best-fitted parameter sets. **(B)** Dose dependent effect of Lapatinib on the expression and phosphorylation of ErbB2 in long-term 10 nM BLU9931-resistant MDA-MB-453 cells. **(C-D)** Treatment of parental or long-term 10 nM BLU9931-resistant MDA-MB-453 cells with BLU9931 (BLU) and Lapatinib (Lap), alone and in combination, at the indicated doses. Cell proliferation was determined by MTS assay. Error bars: mean ± standard error of three biological replicates. * indicates p-value of <0.05, ** < 0.01, *** < 0.001.

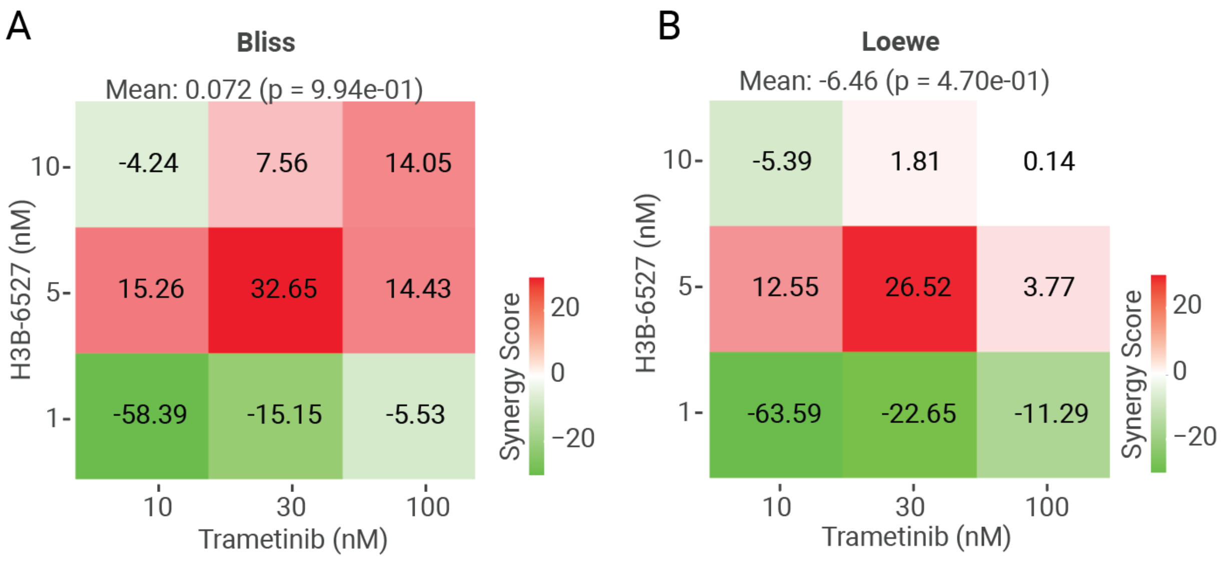


**Figure S22. Quantitative assessment of drug synergism.** Quantitative assessment of drug synergism between H3B-6537 and Trametinib in the MDA-MB-453 cell line using Bliss score (A) and Loewe score (B). To calculate the synergy score, we used SynergyFinder+, an open-source package (47), which can be accessible at www.synergyfinderplus.org.

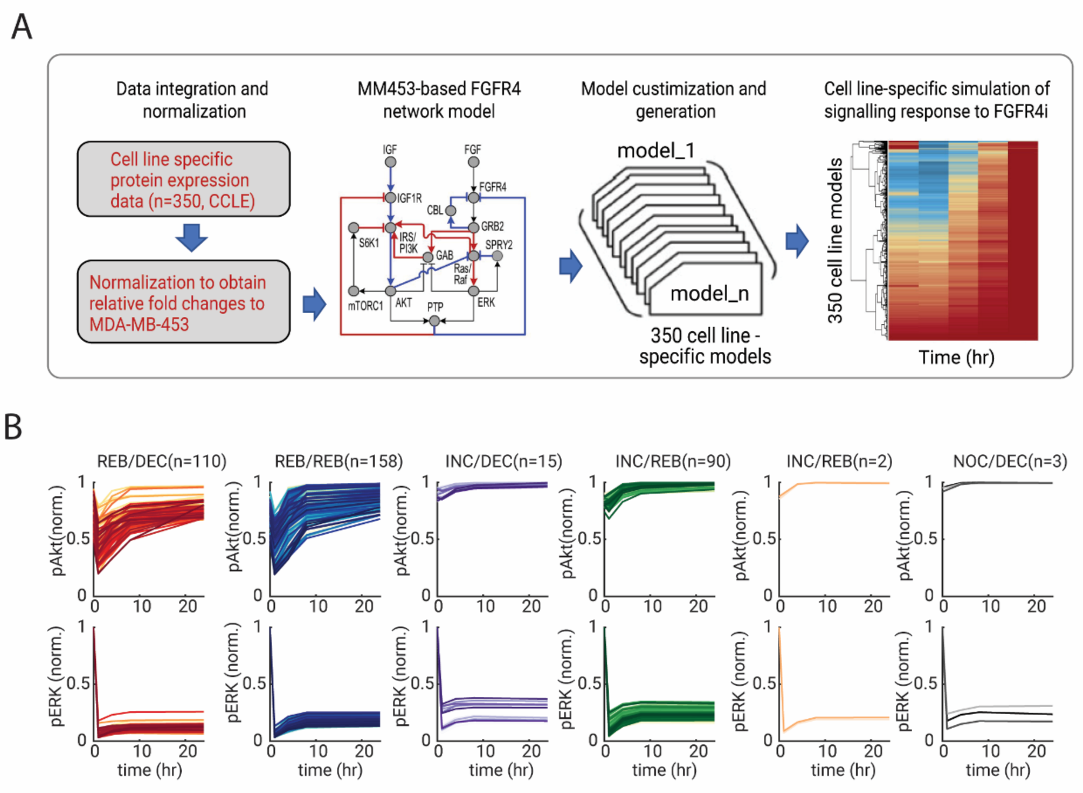


**Figure S23. Cell line-specific simulation of signalling response to FGFR4 inhibition across 350 cancer cell lines. (A)** A flowchart displaying the pipeline of model customisation and cell type specific model generation used in this study. **(B)** Simulated time evolution of AKT and ERK phosphorylation in response to FGFR4 inhibition across 350 cell lines using the cell line specific models generated in (A). The dynamic response curves are overlaid and grouped according to the distinct response pattern subgroups described in the main text.

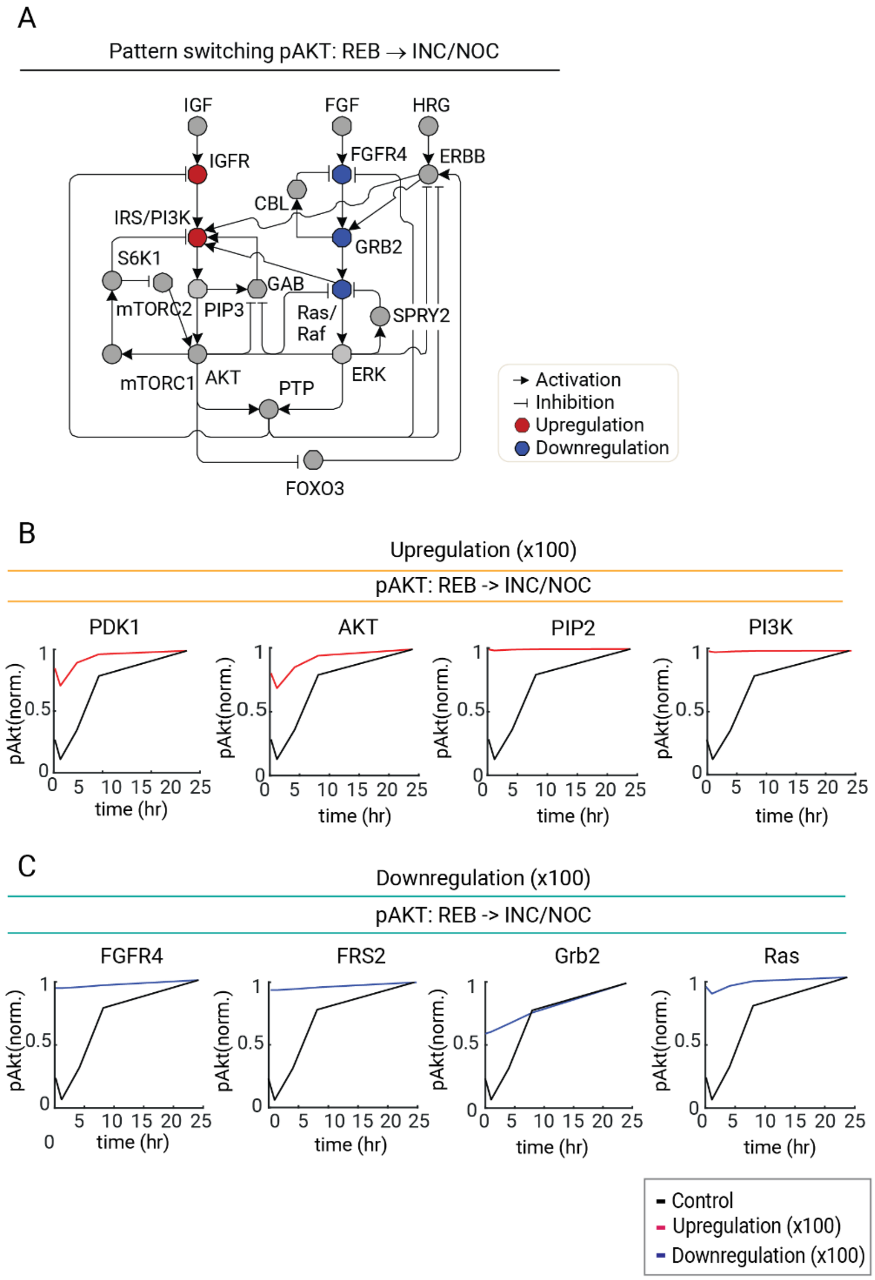


**Figure S24. The perturbation analysis of the FGFR4 signalling Network. (A)** Modelling based identification of network perturbations (i.e. upregulation of the red or downregulation of the blue components) that convert the pAKT response from a REB to an INC/NOC pattern, demonstrating the marked plasticity of network-mediated drug response dynamics. **(B)** Time course simulation demonstrating the conversion of pAKT pattern by perturbation of specific network nodes. Upregulation of the identified network nodes (PDK1, AKT, PIP2, PI3K) or downregulation of the identified network nodes (FGFR4, FRS2, Grb2, Ras) convert pAKT response pattern from REB to INC/NOC.

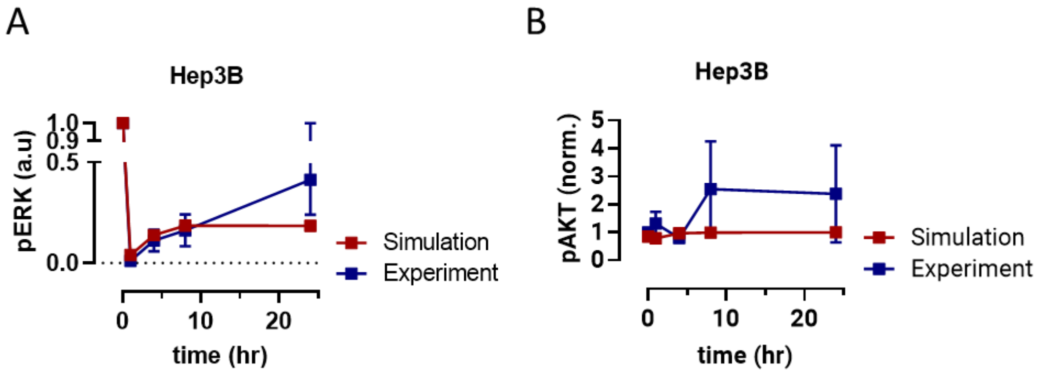


**Figure S25. Effect of the FGFR4 inhibitor H3B-6527 on FGFR4 downstream signalling and proliferation in the Hep3B cell line.** Model prediction and quantification by densitometry of pERK (A) and pAKT dynamics (B) in response to FGFR4 inhibition in Hep3B cells. See Figure 7E for the western blot data. Note that data were first normalised relative to the Tubulin control and the total protein, then phosphorylated proteins were normalised to DMSO control. Error bars: mean ± standard error of three biological replicates.

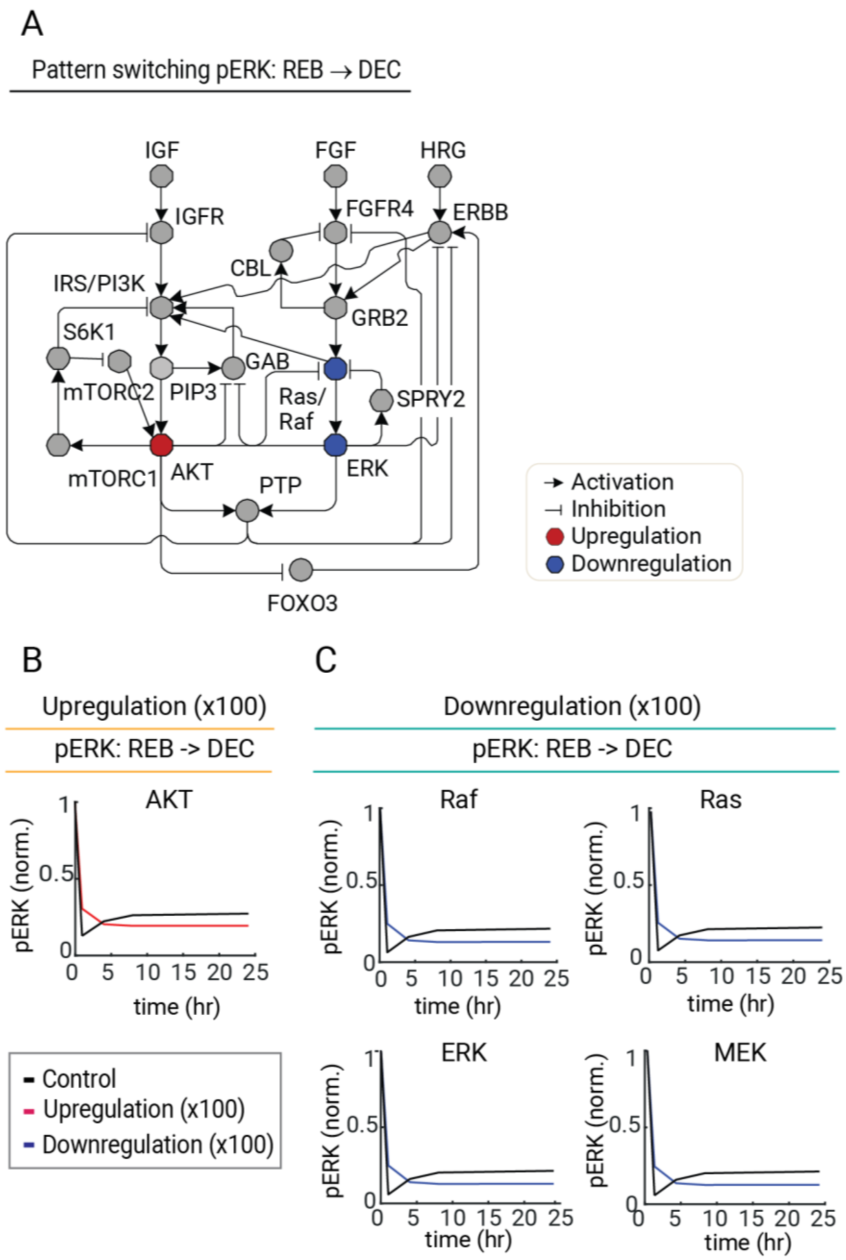


**Figure S26. The perturbation analysis of the FGFR4 signalling Network. (A)** Modelling based identification of network perturbations (i.e. upregulation of the red or downregulation of the blue components) that convert the pERK response from a REB to a DEC pattern, demonstrating the marked plasticity of network-mediated drug response dynamics. **(B)** Time course simulation demonstrating the conversion of pAKT/pERK response pattern by perturbation of specific network nodes. Downregulation of the identified network nodes (Raf, Ras, ERK, MEK) or upregulation of pAKT convert pERK response pattern from REB to DEC.

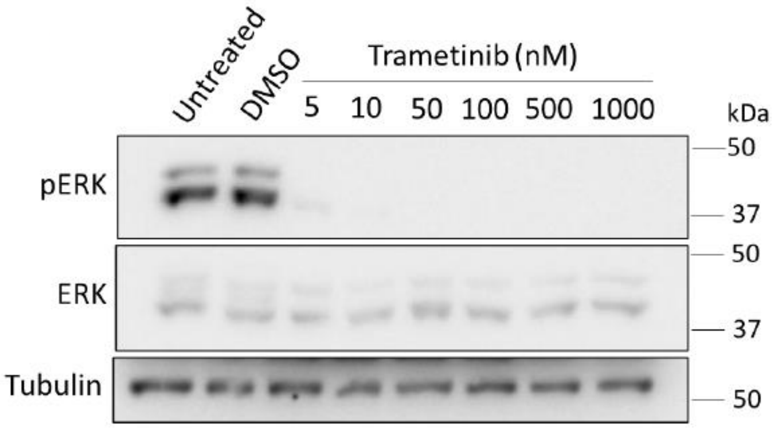


**Figure S27. Dose dependent effect of MEK inhibitor Trametinib on ERK phosphorylation in the Hep3B cell line.** Expression and activation of ERK was characterized after 1 h treatment with the indicated doses.

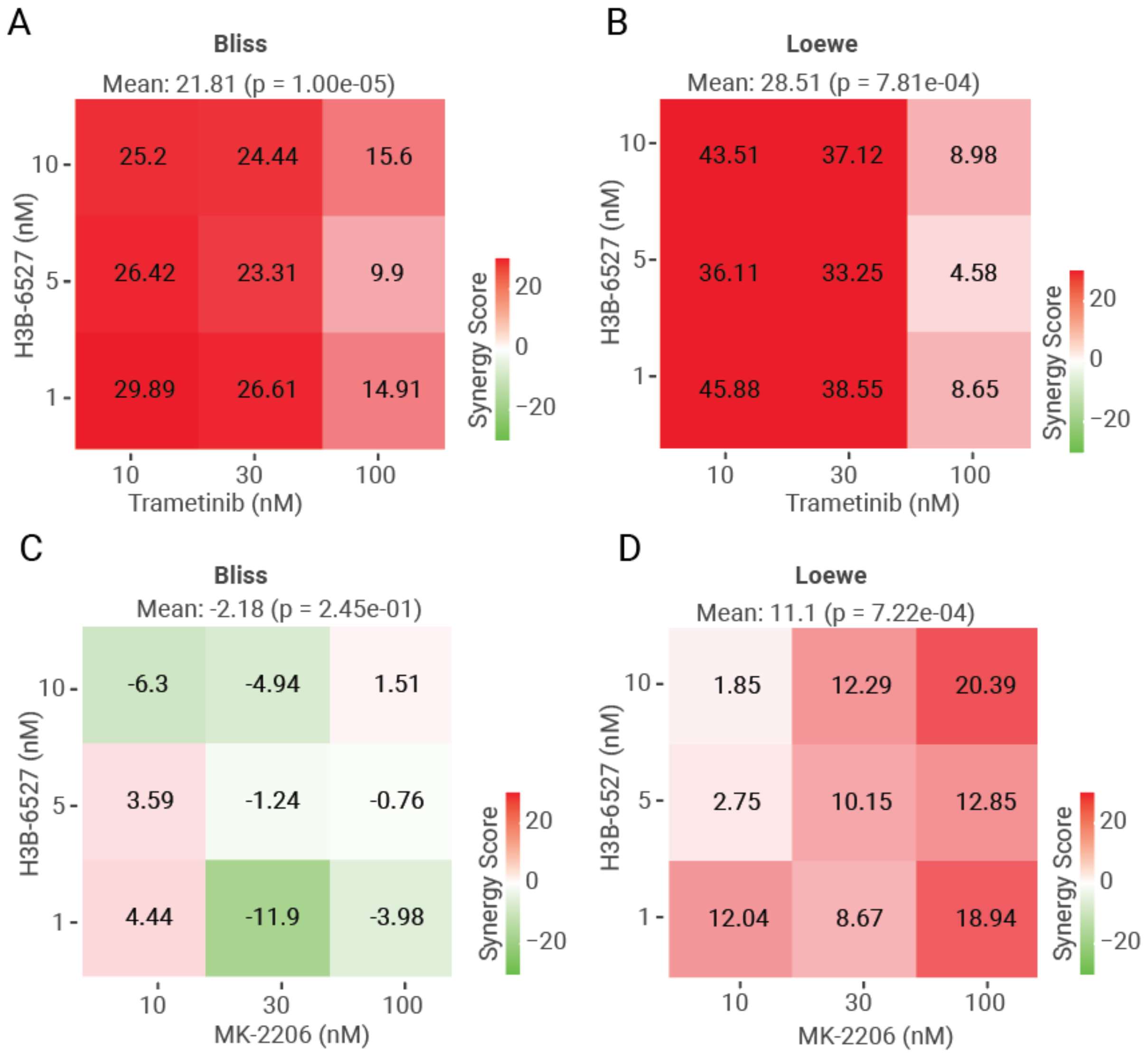


**Figure S28. Quantitative assessment of drug synergism.** Quantitative assessment of drug synergism between H3B-6537 and Trametinib in the Hep3B cell line using Bliss score (A) and Loewe score (B). Quantitative assessment of drug synergism between H3B-6537 and MK-2206 using Bliss (C) and Loewe score (D). To calculate the synergy score, we used SynergyFinder+, an open-source package (47), which can be accessed at accessible at www.synergyfinderplus.org.

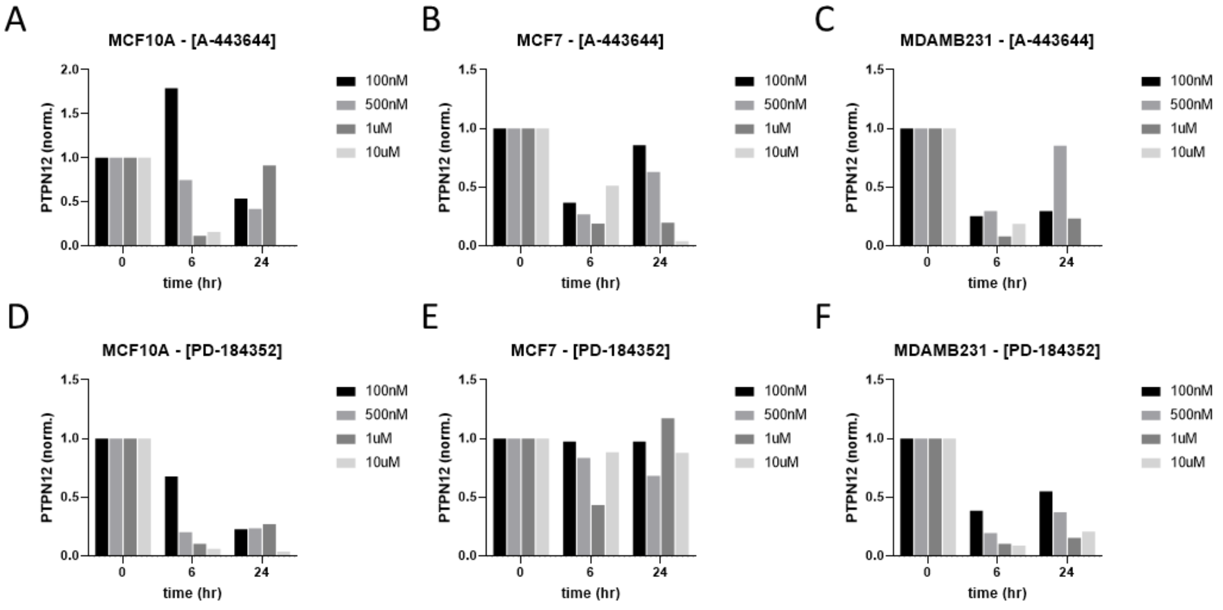


**Figure S29. Gene expression of PTPN12 in response to AKT and MEK inhibition by small molecule drugs.** Gene expression data were downloaded for analysis from the Connectivity Map (CMap) consortium (20,21). A-443644 is a pan-AKT inhibitor. PD-184352 is a MEK inhibitor.

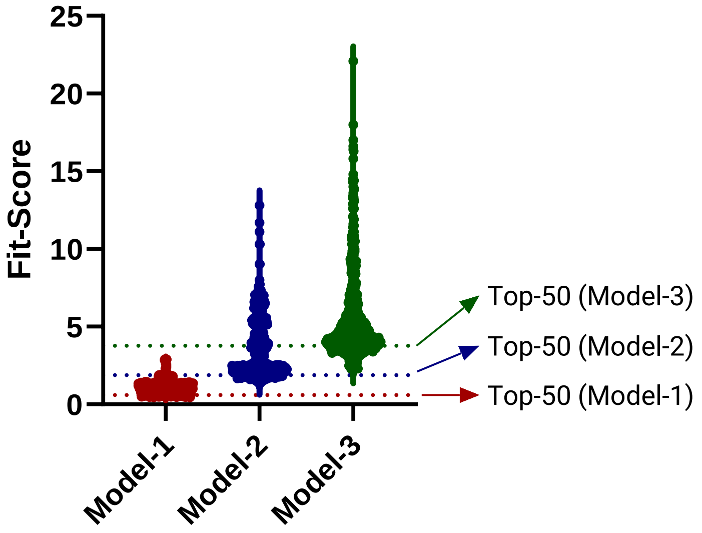


**Figure S30.** Result of the >300 independent GA-based optimization runs for the models of phase-1, -2 and -3. Per model, each GA run produced a fitted parameter set, which were then sorted by their corresponding objective function values (Fit-Score). Top 50 best-fitted parameter sets were selected for further analysis.

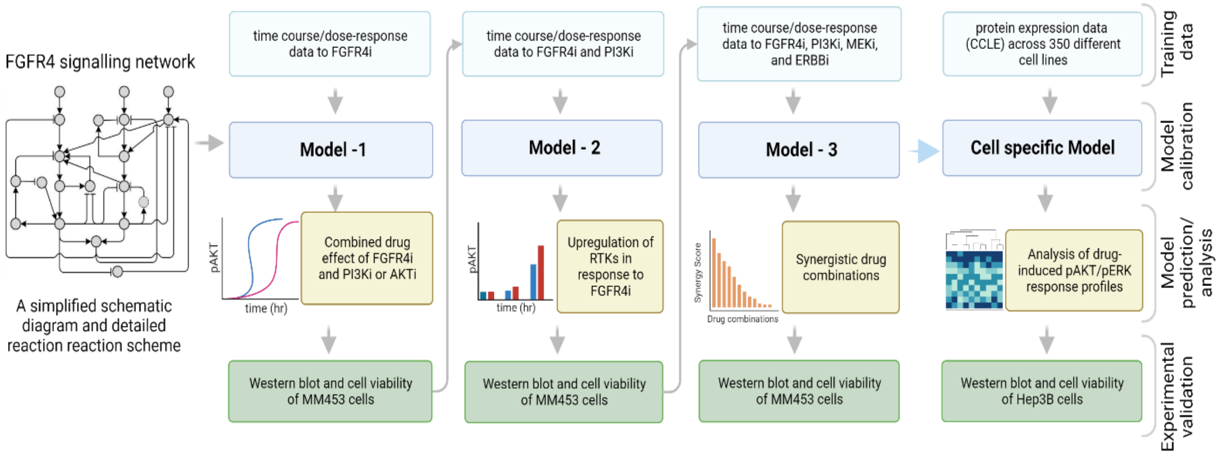


**Figure S31. Workflow of the FGFR4 network modelling and analysis**. The model fitting process contains three sequential phases, where at each phase new data generated to validate new model predictions are then incorporated into the pool of training data and the entire pool was used to recalibrate the model at the next phase. The cell specific model is tailored based on Model-3 and incorporates cancer cell specific protein expression data.

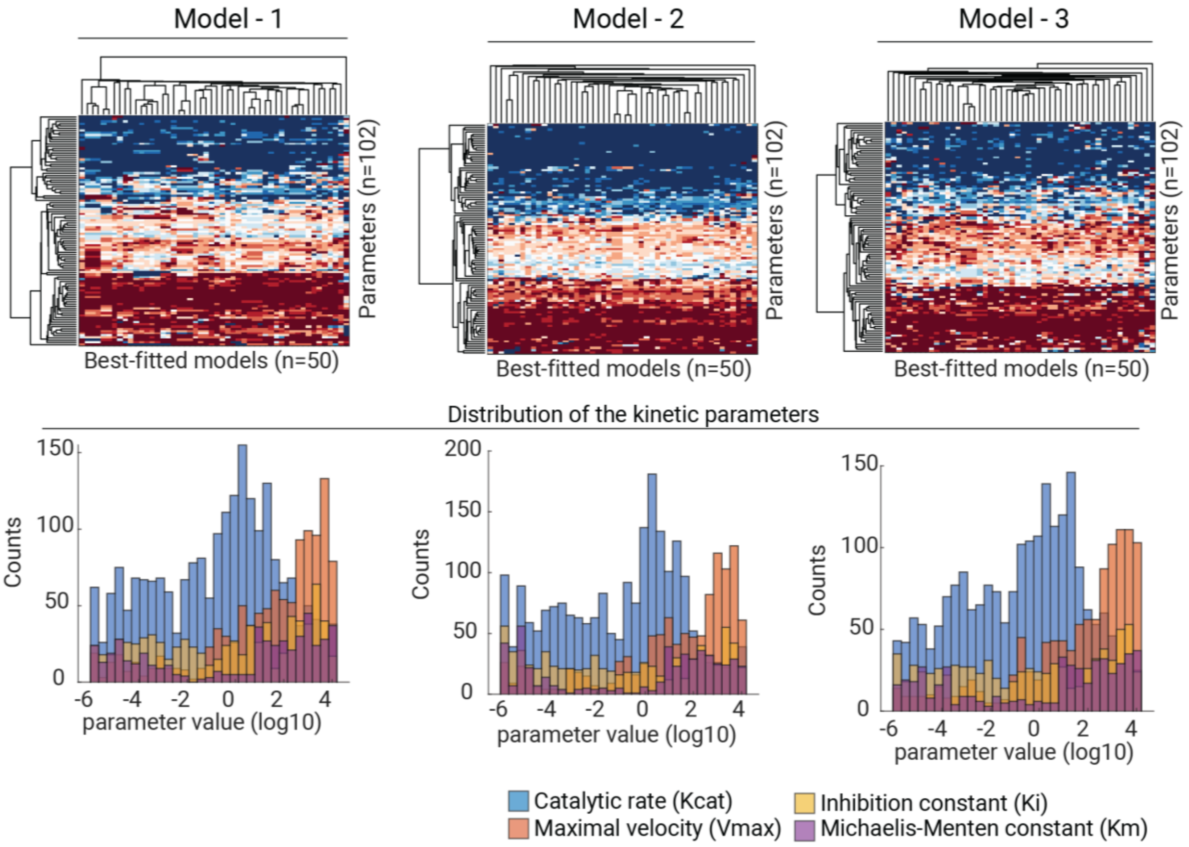


**Figure S32.** Heatmap of the top 50 best-fitted parameter sets (upper panels) and histogram of parameter values of *kcat*, *Vmax*, *Km* and *Ki* (lower panels).

### S3. Supplementary Tables

**Table S1. Reactions and rate equations of the FGFR4 signaling network model.**

| | Reaction | Rate equations |
|---|---|---|
| v01 | IGFR → pIGFR | kc01f * IGF * IGFR - Vm01r * (1 + ki01r*PTP) * pIGFR |
| v02 | FGFR4 → pFGFR4 | kc02f * FGF * FGFR4 / (1 + Ki02f * (FGFR4i^n02 / (Kmf02^n02 + FGFR4i^n02))) - Vm02r * (1 + ki02r1 * PTP) * (1 + ki02r2 * aCbl) * pFGFR4 |
| v03 | ERBB → pERBB | (kc03f * HRG + kc03f2 * FOXO) * ERBB / (1+ ki03f *pERK) / (1 + Ki03f2*(ERBBi/(Kmf03 + ERBBi))) - Vm03r * (1 + ki03r*PTP) * pERBB |
| v04 | IRS → pIRS | kc04f * pIGFR * IRS / (1+ ki04f *pS6K) - Vm04r * pIRS |
| v05 | PI3K → aPI3K | (kc05f1 * pIRS + kc05f2 * (aGAB1 + aGAB2) + kc05f3 * aRas + kc05f4 * pERBB) * PI3K /(1 + Ki06f*(PI3Ki^n06/(Kmf06^n06 + PI3Ki^n06))) - Vm05r * aPI3K |
| v06 | PIP2 → PIP3 | kc06f * aPI3K * PIP2 - kc06r * PTEN * PIP3 |
| v07 | PDK1 → aPDK1 | kc07f * PIP3 * PDK1 - Vm07r * aPDK1 |
| v08 | Akt → pAkt | (kc08f * aPDK1 + kc08f1 * amTORC2) * Akt / (1 + Ki08f*(AKTi/(Kmf08 + AKTi))) - Vm08r * pAkt |
| v09 | mTORC1 → amTORC1 | (kc09f * pAkt + kc09f1 * pERK)* mTORC1 - Vm09r * amTORC1 |
| v10 | S6K → pS6K | kc10f * S6K * amTORC1 - Vm10r * pS6K |
| v11 | FRS2 → pFRS2 | kc11f * pFGFR4 * FRS2 / (1+ki11r*pERK) - Vm11r * pFRS2 * (1+ki11f*aCbl) |
| v12 | Grb2 → aGrb2 | (kc12f1 * pFRS2/(1+ki12f*pSPRY2) + kc12f2 * pERBB + kc12f3 * pIRS) * Grb2 * Grb2 - Vm12r * aGrb2 |
| v13 | Sos → aSos | kc13f * Sos * aGrb2 - Vm13r * aSos |
| v14 | Shp2 → aShp2 | (kc14f1 * aGrb2 + kc14f2 * (aGAB1 + aGAB2)) * Shp2 - Vm14r * aShp2 |
| v15 | Ras → aRas | (kc15f1 * aSos/(1 + ki15f*pERK) + kc15f2 * aShp2)*Ras - Vm15r * aRas |
| v16 | Raf → aRaf | kc16f * Raf * aRas / ((1 + ki16f1*pAkt) * (1 + ki16f2*(SPRY2 + pSPRY2))) - Vm16r * aRaf |
| v17 | MEK → pMEK | kc17f * MEK * aRaf - Vm17r * pMEK |
| v18 | ERK → pERK | kc18f * ERK * pMEK - Vm18r * pERK |
| v19 | GAB1 → aGAB1 | kc19f * (1 + alpha19f * PIP3)* aGrb2 * GAB1 /(1 + ki19f*pERK) - Vm19r * aGAB1 |

| | | |
|---|---|---|
| v20 | GAB2 → aGAB2 | kc20f * (1 + alpha20f * PIP3)* aGrb2 * GAB2 /((1 + ki20f1*pERK) * (1 + ki20f2*pAkt)) - Vm20r * aGAB2 |
| v21 | ∅ → mSPRY2 | vs21 + kc21 * pERK /(Km21 + pERK) |
| v22 | mSPRY2 → ∅ | (vs21 + kc21)/100 * mSPRY2 |
| v23 | ∅ → SPRY2 | kc23 * mSPRY2 / (Km23 + mSPRY2) |
| v24 | SPRY2 →∅ | kc24 * SPRY2 |
| v25 | SPRY2 → pSPRY2 | kc25f * SPRY2 * pFRS2 - kc25r * pSPRY2 |
| v26 | pSPRY2 →∅ | kc26 * pSPRY2 |
| v27 | ∅ → mPTP | vs27 + kc27a * pERK / (Km27a + pERK) + kc27b *pAkt /(Km27b + pAkt) |
| v28 | mPTP → ∅ | (vs27 + kc27a + kc27b)/100 * mPTP |
| v29 | ∅ → PTP | kc29 * mPTP / (Km29 + mPTP) |
| v30 | PTP → ∅ | kc29/100 * PTP |
| v31 | Cbl → aCbl | kc31f * aGrb2 * Cbl - Vm31r * aCbl |
| v32 | FOXO → pFOXO | kc32f * pAkt * FOXO - Vm32r * pFOXO |
| v33 | mTORC2 → amTORC2 | kc33f * PIP3 * mTORC2 /(1 + ki33f*pS6K) - Vm33r * amTORC2 |

**Table S2. Ordinary differential equations of the integrated FGFR4 model.** The reaction rates are given in Supplementary Table S1.

| **Left-hand Sides** | **Right-hand Sides** |
|---|---|
| d[IGFR]/dt | -v01 |
| d[pIGFR]/dt | +v01 |
| d[FGFR4]/dt | -v02 |
| d[pFGFR4]/dt | +v02 |
| d[ERBB]/dt | -v03 |
| d[pERBB]/dt | +v03 |
| d[IRS]/dt | -v04 |
| d[pIRS]/dt | +v04 |
| d[PI3K]/dt | -v05 |
| d[aPI3K]/dt | +v05 |
| d[PIP2]/dt | -v06 |
| d[PIP3]/dt | +v06 |
| d[FRS2]/dt | -v11 |
| d[pFRS2]/dt | +v11 |

| | |
|---|---|
| d[Grb2]/dt | -v12 |
| d[aGrb2]/dt | +v12 |
| d[Akt]/dt | -v08 |
| d[pAkt]/dt | +v08 |
| d[PDK1]/dt | -v07 |
| d[aPDK1]/dt | +v07 |
| d[mTORC1]/dt | -v09 |
| d[amTORC1]/dt | +v09 |
| d[S6K]/dt | -v10 |
| d[pS6K]/dt | +v10 |
| d[Sos]/dt | -v13 |
| d[aSos]/dt | +v13 |
| d[Shp2]/dt | -v14 |
| d[aShp2]/dt | +v14 |
| d[Ras]/dt | -v15 |
| d[aRas]/dt | +v15 |
| d[Raf]/dt | -v16 |
| d[aRaf]/dt | +v16 |
| d[MEK]/dt | -v17 |
| d[pMEK]/dt | +v17 |
| d[ERK]/dt | -v18 |
| d[pERK]/dt | +v18 |
| d[GAB1]/dt | -v19 |
| d[aGAB1]/dt | +v19 |
| d[GAB2]/dt | -v20 |
| d[aGAB2]/dt | +v20 |
| d[mSPRY2]/dt | +v21-v22 |
| d[SPRY2]/dt | +v23-v24-v25 |
| d[pSPRY2]/dt | +v25-v26 |
| d[mPTP]/dt | +v27-v28 |
| d[PTP]/dt | +v29-v30 |
| d[Cbl]/dt | -v31 |
| d[aCbl]/dt | +v31 |
| d[FOXO]/dt | - v32 |
| d[pFOXO]/dt | +v32 |
| d[mTORC2]/dt | -v33 |
| d[amTORC2]/dt | +v33 |

**Table S3. FGFR4 kinase domain primers.**

| Primer | Primer sequence (5’ to 3’) | Tm (°C) |
|---|---|---|
| 1F | AGATGCTCAAAGACAACGCC | 58 |
| 1R | AGATACTGCATGCCTCGGG | 59 |
| 2F | CACTGTGCAGAAGCTCTCCC | 59 |
| 2R | AAGGTCGAGCACTGTGTCAG | 59 |
| 3F | TCTCGACCCACTATGGGAGT | 59 |
| 3R | TTGTCCTCAGTCACCAGCAC | 59 |
| 4F | GCCGGCCTCGTGAGTCTA | 59 |
| 4R | TACACTTCCGGGACTCCAGAT | 59 |
| 5F | CAGAAGCTCTCCCGCTTCC | 59 |
| 5R | CCGAGCAGAACCCTGACATT | 59 |

**Table S4. Constituent readouts making up the ICV function**, along with inferred relative weights from maximal drug effect data by targeted agents.

| Model readouts | Targeting drugs | Maximal drug effect | Contributing Weights | Source databases |
|---|---|---|---|---|
| pAKT | MK-2206 | 0.024 | 0.976 | CTD |
| pERK | BVD-523 | 0.59 | 0.41 | GDSC2 |
| pS6K | PF-4708671 | 0.049 | 0.921 | GDSC1 |

**S4. Supplementary Data**

**Data S1. The SBML model file for the integrated FGFR4 signalling network.** This is provided in a separate xml file.

**Data S2. Estimated best-fitted parameter sets (Model-1)**. This is provided in a separate excel file.

**Data S3. Estimated best-fitted parameter sets (Model-2)**. This is provided in a separate excel file.

**Data S4. Estimated best-fitted parameter sets (Model-3)**. This is provided in a separate excel file.
